# Site-specific combinatorial ubiquitination drives the targeted degradation of plasma membrane proteins

**DOI:** 10.64898/2026.05.23.727405

**Authors:** Niclas Olsson, Aleksandr Gaun, Sammy Villa, Katrina Jackson, Mark E. Fitzgerald, Fiona E. McAllister, David Stokoe, Aaron H. Nile, Kirill Bersuker

## Abstract

Cullin-RING E3 ligases (CRLs) ubiquitinate their substrates by recruiting them to complexes containing ubiquitin-conjugating enzymes (UCEs). Ubiquitination proceeds through the transfer of ubiquitin from UCEs to substrate lysines, but the specificity of lysine ubiquitination has not been widely investigated in cells. In this study, we use global ubiquitination profiling to identify sites in the kinase domains of receptor tyrosine kinases (RTKs), EGFR and Her2, that undergo CRL-mediated ubiquitination in cells treated with heterobifunctional degrader molecules. We find that VHL- and Cereblon-dependent degraders trigger the ubiquitination of a common subset of accessible lysines. Using mutagenesis, we show that the number of available ubiquitination sites specifies the extent of RTK degradation. Furthermore, introduction of non-native sites mostly fails to rescue clearance of lysine-deficient RTK mutants. These results highlight the specificity with which UCEs ubiquitinate their substrates and suggest that the ubiquitination of multiple sites governs the efficacy of degraders targeting plasma membrane proteins.

## Introduction

Protein degradation is governed by the post-translational covalent modification of primary amines with ubiquitin, a signal that targets its substrates for proteolysis by proteasomes and lysosomal proteases^1^ . The multitude of ways by which ubiquitin can be conjugated to proteins, such as a monomer or chain of monomers appended to one or more lysines within the modified substrate or to ubiquitin itself, endows ubiquitin with the ability to trigger degradation through distinct pathways and function as a key regulator of most other cellular processes. Furthermore, the dynamic nature of ubiquitin conjugation, governed by the opposing actions of E3 ligases and deubiquitinating (DUB) enzymes that add and remove ubiquitin, respectively, gives cells spatial and temporal control over ubiquitin signaling. Ubiquitin conjugation is controlled in part by the activity of E3 ligase complexes, whose function is to recognize substrates, bind ubiquitin, and position substrates in spatial configurations that facilitate ubiquitin transfer. For Cullin-RING E3 ligases (CRLs), these activities are executed by substrate adaptors that position the substrate within the CRL complex, and E2 ubiquitin-conjugating enzymes (UCEs) that covalently bind ubiquitin^2^. Ubiquitination occurs when the relative orientations of the substrate and the ubiquitin-bound UCE create conditions that are energetically favorable for ubiquitin transfer to the substrate^3,4^. How CRL complexes spatially position thousands of distinct substrates to facilitate their ubiquitination by a much more limited repertoire of UCEs is central to understanding the ubiquitin system.

The reconstitution of CRL complexes *in vitro* and the elucidation of their structures have shed light on specific interactions that promote ubiquitin transfer. A recent study that reconstituted a Cullin 2 (Cul2)-based CRL complex containing the ubiquitin-bound UCE UBE2R2 and the substrate adaptor FEM1C highlighted the importance of the UCE-substrate adaptor interaction during the ubiquitination of a model peptide substrate^3^. To investigate the ubiquitination of a larger CRL substrate, the authors solved a structure using a heterobifunctional degrader that recruits the BD2 domain of BRD4 to the Cul2 CRL adaptor VHL. A combination of mass spectrometry and mutagenesis was used to map ubiquitinated lysines in BRD4-BD2 and identify a key lysine necessary for ubiquitination by the UCE. Another study that reconstituted the same complex identified a ubiquitination zone consisting of multiple lysines in BRD4-BD2 within a region of the substrate spatially oriented towards the UCE^5^. These studies demonstrated the utility of using degraders to structurally model interactions of CRLs with their substrates and illustrated how the positioning of UCEs, substrate adaptors, and substrates within CRLs promotes ubiquitin transfer.

The insights gained from structural and activity analyses of reconstituted CRL complexes have motivated questions as to how CRLs ubiquitinate their substrates within the compartmentalized interiors of intact cells. The dynamic nature of ubiquitination increases the difficulty of capturing ubiquitinated species, which are often present in low abundance without perturbation of degradation machinery and DUBs^6,7^. Degradation systems that use small molecules to promote the ubiquitin-dependent degradation of specific tags can, in principle, be used to study ubiquitination events, but they are limited to a handful of substrates for which ligands have been identified^8^. These limitations can be overcome by using degraders to ubiquitinate a substrate of interest, provided that bifunctional molecules for a given substrate adaptor and target pair are available. The ability to trigger recruitment of CRLs to membrane-associated proteins also presents an opportunity to understand how ubiquitination signals the degradation of distinct classes of substrates^9^.

In this study we use bifunctional degraders targeting the kinase domains of plasma membrane-anchored receptor tyrosine kinases (RTKs) and global ubiquitin proteomics to profile CRL-dependent ubiquitination in intact cells. We use these findings to determine the extent to which combinatorial ubiquitination signals degradation of plasma membrane-associated proteins by the endolysosomal system.

## Results

### Development of a Cereblon-dependent Her2 heterobifunctional degrader

To investigate the ubiquitination of a model CRL substrate in cells, we used a series of heterobifunctional degrader molecules to recruit substrate adaptors to the kinase domains (KDs) of oncogenic RTKs^10^. Lapatinib-VHL (Lap-VHL) binds to the ATP-binding pocket of Her2 and EGFR and to the CRL substrate adaptor VHL, which functions as part of the Cullin 2 (Cul2) CRL (Figure S1A)^11,12^. To determine how replacing the KD binding group impacts ubiquitination, we used the Her2-specific KD binder, Tucatinib-VHL (Tuc-VHL)^12^ (Figure S1B). Additionally, we modified the identity of the ubiquitinating CRL by synthesizing Tucatinib-Cereblon (Tuc-CRBN), a degrader that consists of a CRBN-binding pyrimidine-glutarimide scaffold linked to Tucatinib through a bipiperidinyl linker (Figure S1C and Supplemental Methods). This compound was designed to trigger the degradation of Her2 by forming a complex between its KD and the Cullin 4 (Cul4) CRL substrate adaptor, Cereblon^13^. An overnight treatment of Her2-amplified SK-BR-3 cells with Tuc-CRBN resulted in the degradation of Her2 with an IC50 of 4.5 nM and a Dmax of ∼50%. In contrast, treatment with a control methylated compound unable to bind CRBN (Tuc-CRBN_Inactive_) did not decrease Her2 levels^14^ (Figures 1A,B and S1C). However, a shorter 2-hour treatment with either active or inactive Tuc-CRBN reduced Her2 phosphorylation, indicating that both compounds bind to and inhibit the kinase activity of the Her2 KD with similar potency (Figure 1C).

**Figure 1.**
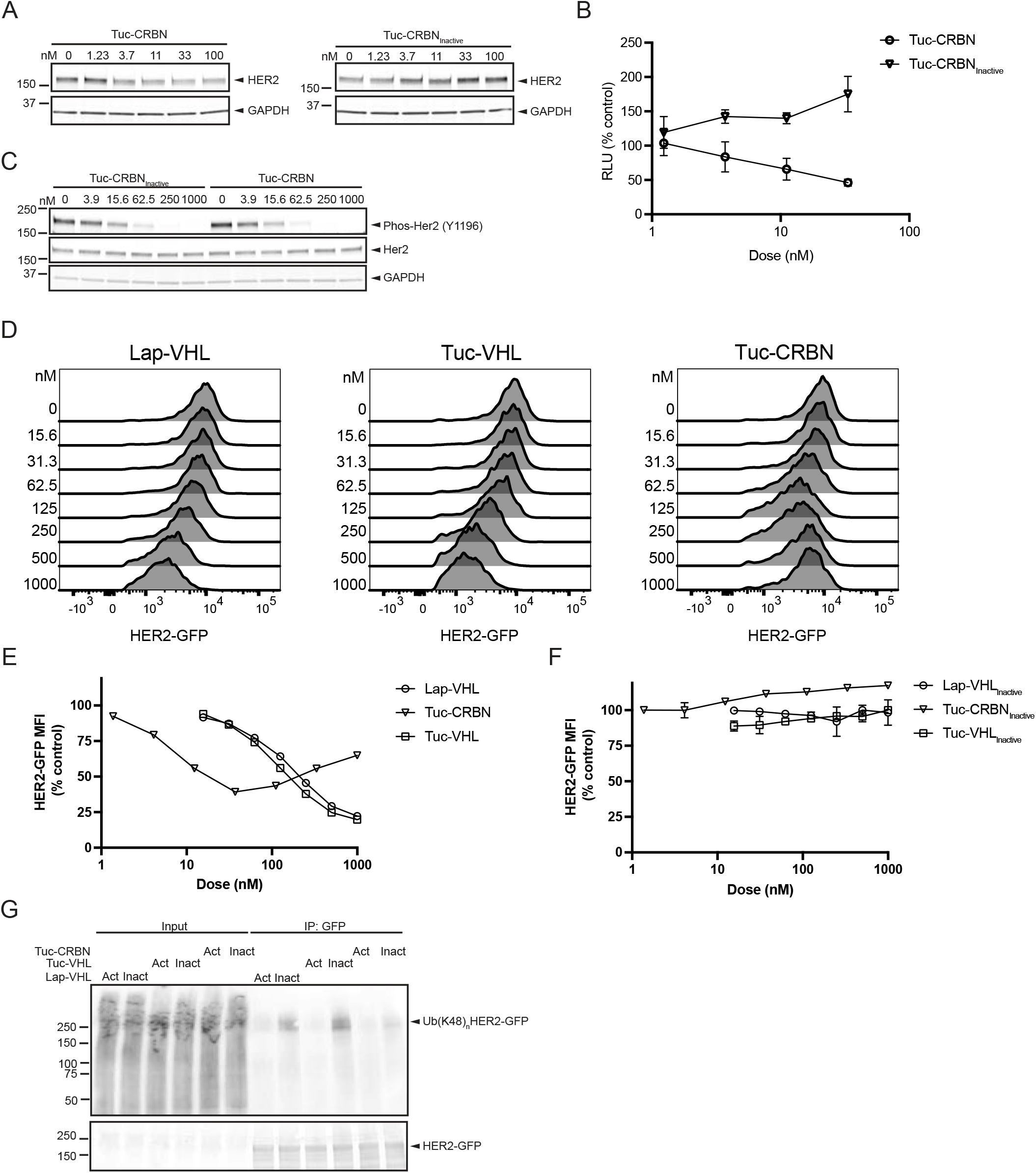
Characterization of Tuc-CRBN activity in cells expressing Her2-GFP. A, B. Western blot analysis of Her2 levels in lysates from SK-BR-3 cells treated with active or inactive Tuc-CRBN for 20 hrs. (A) shows a blot representative of three biological replicates and (B) shows quantification of Her2 levels as mean ± SD. C. Western blot analysis of phosphorylated Her2 levels in lysates from cells treated for 2 hrs with degraders. The western blot is representative of two biological replicates. D-F. Flow cytometric analysis of Her2-GFP levels in cells treated with the indicated degraders. The mean Her2-GFP fluorescence intensity of the cell population after a 20 hr treatment with active (E) or inactive (F) degraders is shown as mean ± SD of 3 technical replicates. G. Western blot analysis of Her2-GFP immuno-purifications from cells treated with active and inactive Lap-VHL (1 μM), Tuc-VHL (1 μM) or Tuc-CRBN (40 nM) for 2 hrs. An Anti-GFP antibody was used to detect Her2-GFP. The blot shows the results of two biological replicates.

### Degrader molecules trigger site-specific ubiquitination of RTK kinase domains

We generated a transgenic SK-BR-3 cell line stably expressing Her2 C-terminally tagged with green fluorescent protein (Her2-GFP) to track the expression levels and ubiquitination status of Her2. Treatment of these cells with degraders resulted in a dose-dependent decrease in Her2-GFP levels (Figure 1D). Tuc-CRBN decreased both Her2-GFP and endogenous Her2 levels to a similar extent but triggered a hook-dependent rebound in Her2-GFP at higher doses, an effect that has been observed when using CRBN-dependent degraders against other targets^15^ (Figures 1D and 1E). As expected, inactive degraders did not reduce Her2-GFP levels (Figure 1F and S1A–C). To detect ubiquitinated Her2, we immunopurified Her2-GFP from cells treated for 2 hrs with degraders and probed for the presence of lysine K48-linked ubiquitin chains, the linkage type assembled by VHL- and CRBN-associated CRLs^4^. Active degraders, unlike the inactive control molecules, increased the levels of K48 ubiquitin chains associated with immunopurified Her2-GFP. This indicates that at shorter time scales, degraders induce ubiquitination prior to degradation of Her2 (Figure 1G).

To identify RTK residues that undergo degrader-dependent ubiquitination, we used remnant motif (K-ε-GG) proteomics to detect ubiquitinated peptides^16^. With this technique, ubiquitinated peptides are enriched using an antibody directed against lysine conjugated to the C-terminal glycine residues (K-ε-GG) of ubiquitin. K-ε-GG peptides immunopurified from cells treated with active and inactive degraders or vehicle control were multiplexed using tandem mass tags (TMT), which were resolved to determine the identity and relative abundance of ubiquitinated sites (Figure S2A and Supplemental Data S1). Consistent with the binding of lapatinib to the KDs of Her2 and EGFR, we detected an increase in ubiquitination of both RTKs after treating cells with Lap-VHL (Figure S2B). To compare these sites, we mapped them onto the crystal structures of EGFR KD bound to lapatinib and that of the Her2 KD bound to the Her2-specific inhibitor TAK285^17,18^. The EGFR K-ε-GG sites exhibiting the greatest fold changes, defined as at least a 1.5 fold increase in abundance (Figure S2E), were positioned on the KD face opposite to the predicted location of the Lap-VHL linker (Figures 2A and 2B), suggesting that the KD face at the VHL-EGFR interface is inaccessible to ubiquitination. K762, the most significantly ubiquitinated site in Her2 (Figures 2C and 2D), shares a similar spatial orientation with K860 in EGFR, whereas ubiquitination of homologous lysine in EGFR (K750) was not detected (Figures 2E and S2E). Conversely, Her2 contains an arginine at the position corresponding to EGFR K860. K716 and K854 in Her2, homologous to K708 and K846 in EGFR, respectively, were more ubiquitinated following Lap-VHL treatment (Figures 2E and S2E). Ubiquitination of K1096, which is located within the unstructured C-terminal portion of the Her2 KD, increased, while several other ubiquitinated sites were not impacted (Figure 2C).

**Figure 2.**
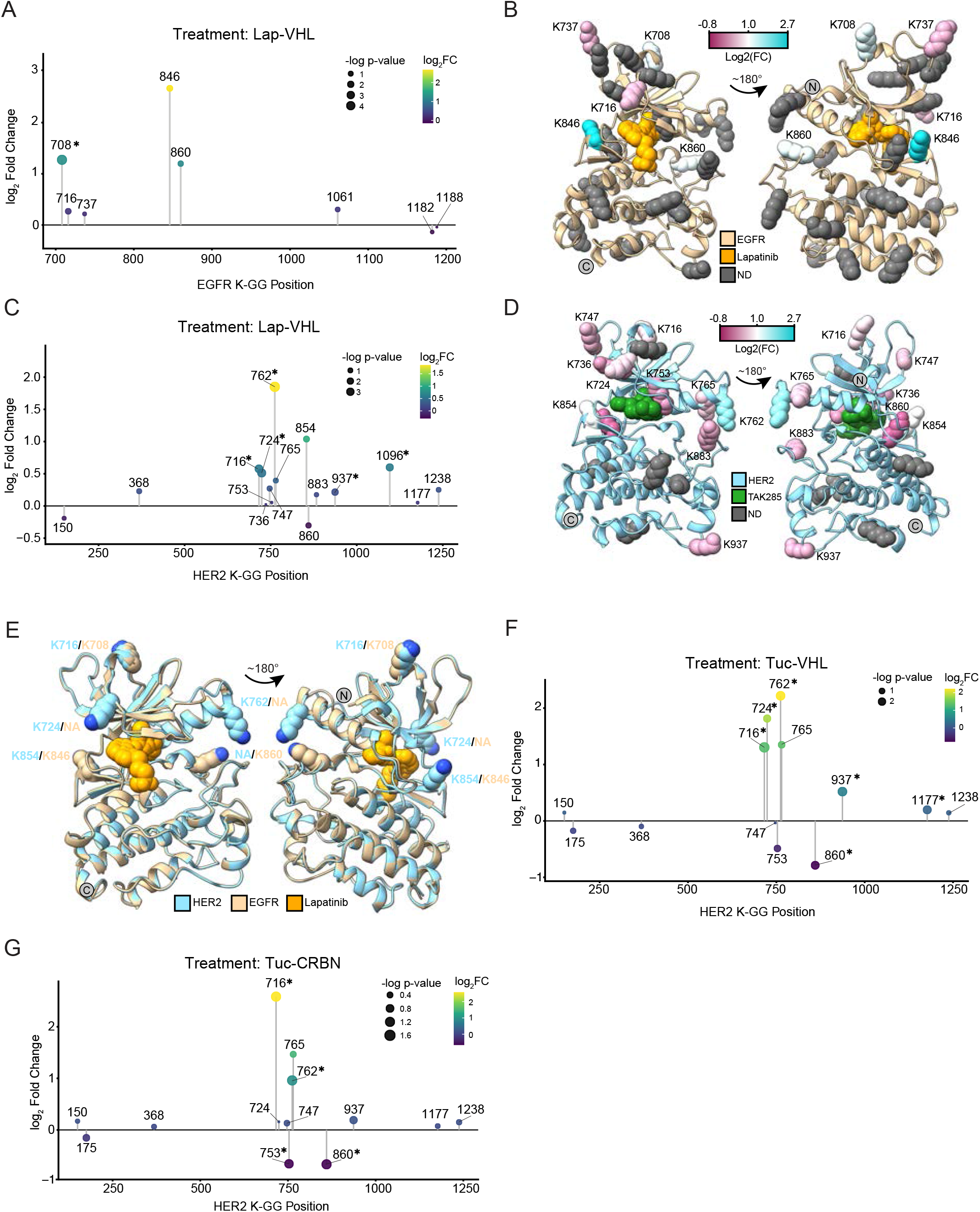
Degraders trigger ubiquitination of specific lysines in RTK kinase domains. A. Fold change in K-GG site abundance in EGFR for cells treated with 1 μM Lap-VHL. B. Mapping of ubiquitinated sites onto the crystal structure of EGFR bound to lapatinib. C. Fold change in K-GG site abundance in Her2 for cells treated with 1 μM Lap-VHL for 3 hrs. D. Mapping of ubiquitinated sites onto the crystal structure of Her2 bound to TAK285. E. Overlay of the EGFR and Her2 crystal structures showing ubiquitinated sites with the greatest fold changes for both KDs. F,G. Fold change in K-GG site abundance for cells treated for 3 hrs with 1 μM Tuc-VHL (F), or 40 nM Tuc-CRBN (G). *p < 0.05 by MS Test. Figures A,C show the results of two biological replicates and Figures F,G show the results of three biological replicates. For statistical analysis of K-GG site abundance, inactive degrader-treated samples were combined with DMSO-treated samples as the comparator group to the active degrader-treated samples.

Treatment with Tuc-VHL impacted similar sites on Her2, suggesting that the Her2-VHL complex formed by Lap-VHL and Tuc-VHL similarly orient the Her2 KD with respect to UCEs (Figures 2F and S2F). While the relative abundance of ubiquitinated K937 and K765 was higher than that observed with Lap-VHL, ubiquitinated K854 peptide was not detected (Figure 2F). Interestingly, Tuc-CRBN induced the ubiquitination of similar sites as Tuc-VHL, with a preference for K716 over K762 (Figures 2F and S2G). This finding was unexpected, given that the CRBN-containing CRL contains distinct adaptors and UCEs^19,20^. Furthermore, the ubiquitination of K753 and K860 decreased in response to both Tuc-VHL and Tuc-CRBN (Figures 2F and 2G). As K753 is required for the KD to bind ATP, degrader binding to the ATP-binding pocket may interfere with constitutive ubiquitination of these lysines^21^. Together, these results indicate that while several lysines are ubiquitinated in inactive degrader and vehicle-treated cells, only a subset possesses the spatial orientation required for further ubiquitination by CRL-associated UCEs.

### Ubiquitination at multiple sites promotes Her2 degradation

Having previously demonstrated that degraders induce RTK clearance by the endolysosomal pathway^12^, we next sought to determine how ubiquitination at specific sites impacts degradation efficiency. To do this, we generated Her2-GFP constructs containing lysine-to-arginine (Lys-Arg) mutations. We focused on a quadruple mutant (Her2-GFP 4KR) containing mutations in K716, K762, K765, and K854, as these sites showed the greatest fold-change in K-ε-GG abundance across all degraders. To confirm that these mutations do not disrupt binding to the Her2 KD, we assessed if degraders can inhibit KD phosphorylation activity. Phosphorylation of the 4KR mutant, wild-type Her2-GFP, and untagged Her2 was inhibited to a comparable extent, confirming that the Lys-Arg mutations do not compromise degrader binding (Figures S3A–C). In contrast, active degraders induced ubiquitination of wild-type Her2-GFP but not the 4KR mutant (Figures 3A–C). Consistent with the lack of ubiquitination, dose-dependent degradation of the 4KR mutant was significantly reduced relative to wild type after Lap-VHL or Tuc-VHL treatment and blocked in response to Tuc-CRBN treatment (Figures 3D–F and 3G–K, *magenta vs. black traces*).

**Figure 3.**
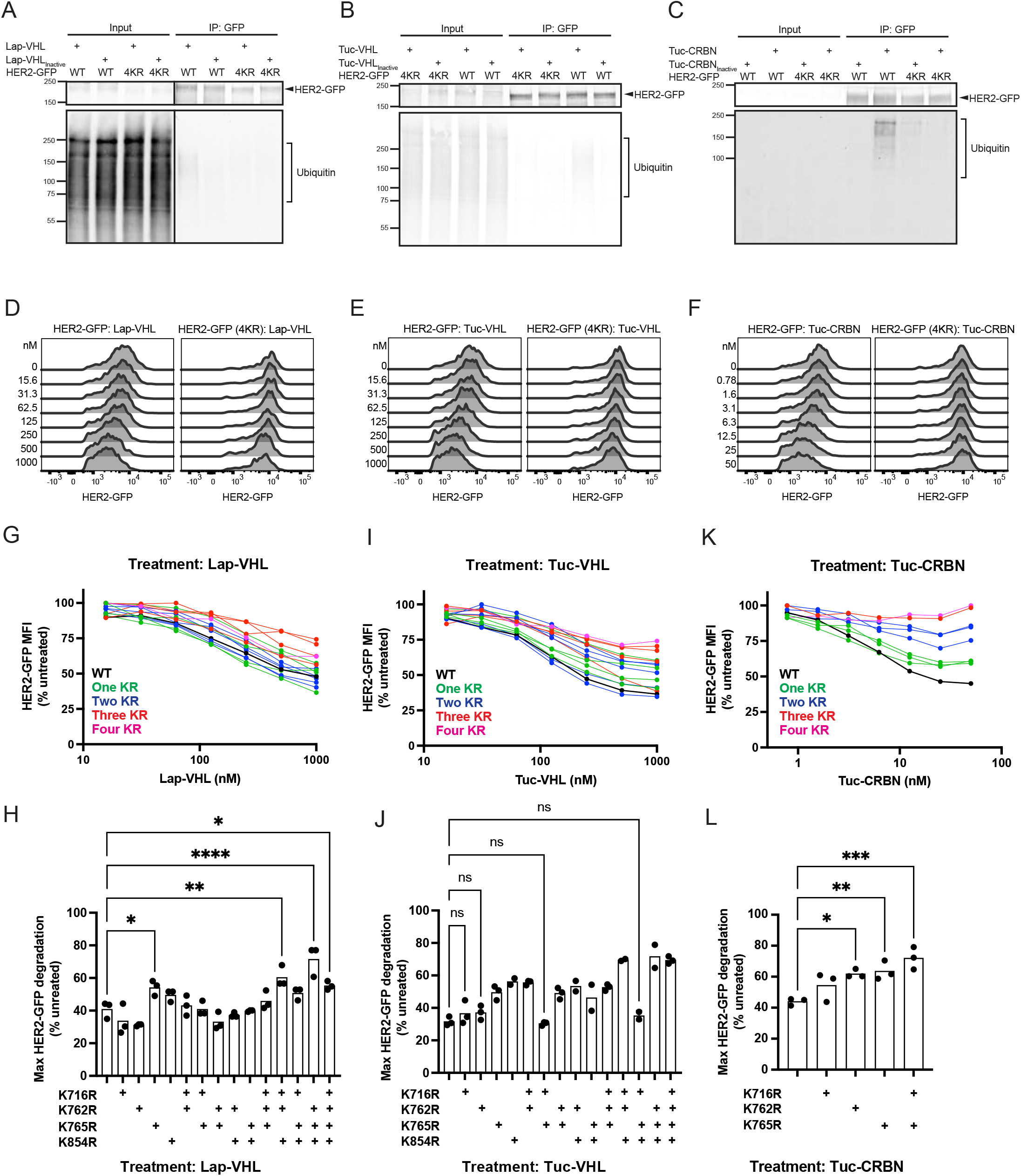
Combinatorial lysine ubiquitination specifies the extent of Her2 degradation. A-C. Western blot analysis of Her2-GFP immunopurifications from cells treated with Lap-VHL (1 μM) (A), Tuc-VHL (1 μM) (B) or Tuc-CRBN (40 nM) (C) for 2 hrs. An anti-GFP antibody was used to detect Her2-GFP. The blots show the results of single experiments. D-F. Flow cytometric analysis of Her2-GFP or Her2 (4KR)-GFP levels in cells treated for 20 hrs with the indicated degraders. 4KR indicates the Her2 KD that contains the K716R, K762R, K765R, K854R mutations. G-K. Quantification of mean Her2-GFP fluorescence of cells expressing WT Her2-GFP, or Her2-GFP mutants containing one to four Lys-Arg mutations. Each data point is the mean of three technical replicates, and each panel is representative of three biological replicates, unless indicated in the quantification below. H-L. Quantification of maximum Her2-GFP degradation across all Her2-GFP mutants and degrader treatments from (G-K). Each data point represents an individual biological replicate. For H,L, significant differences between WT and mutant Her2-GFP lines are indicated; *p < 0.05, **p < 0.01, ***p < 0.001, ****, p < 0.0001. For J, all differences were significant except where indicated. ns, p>0.05. WT vs. K765R/K854R, *p < 0.05. WT vs. K762R/K765R, **p < 0.01. WT vs. K762/K854, WT vs. K716R, K762R, K765R, **p < 0.001. WT vs. all other mutants, ****p < 0.0001. P values were calculated using ordinary one-way ANOVA corrected with Dunnett’s multiple comparisons test.

To determine how each ubiquitination site contributes to Her2 degradation, we expressed Her2-GFP mutants containing between one and four Lys-Arg mutations. In general, increasing the number of mutations attenuated degradation, with maximum inhibition occurring after mutation of three or more lysines (Figures 3G–L). The relationship between site availability and degradability was most evident with Tuc-CRBN treatment, where mutation of any single ubiquitination site resulted in partial inhibition (Figures 3K and 3L). Importantly, these mutations did not impact degrader potency as measured by DC_50_ (Figures S3D–F), in agreement with kinase activity data showing that degrader binding to the Her2 KD was not impacted. Unexpectedly, mutation of K724 both increased degradation beyond that of wild type and blocked the inhibitory effects of other mutations, suggesting that this change may trigger utilization of alternative ubiquitination sites or impacts the intrinsic stability of Her2 (Figure S3G). These findings demonstrate that multiple sites, once ubiquitinated, function combinatorially to drive degradation.

Given the specificity of lysine ubiquitination, we asked if introduction of non-native ubiquitination sites would restore degradation of mutant Her2-GFP. We tested this possibility by introducing individual arginine-to-lysine (Arg-Lys) mutations into the 4KR mutant at positions proximal to the ubiquitinated lysines (Figure 4A). All Arg-Lys mutations failed to rescue mutant Her2-GFP degradation following treatment with Lap-VHL (Figure 4B). However, Tuc-VHL and Tuc-CRBN-dependent degradation was partially restored by the R896K mutation (Figures 4C and 4D). Remarkably, mutation of the neighboring arginine (R897K) did not have this effect, potentially due to the less favorable spatial orientation of that specific lysine. These results indicate that lysines proximal to known ubiquitinated sites are not necessarily targeted, thus further highlighting the specificity by which CRLs ubiquitinate their substrates.

**Figure 4.**
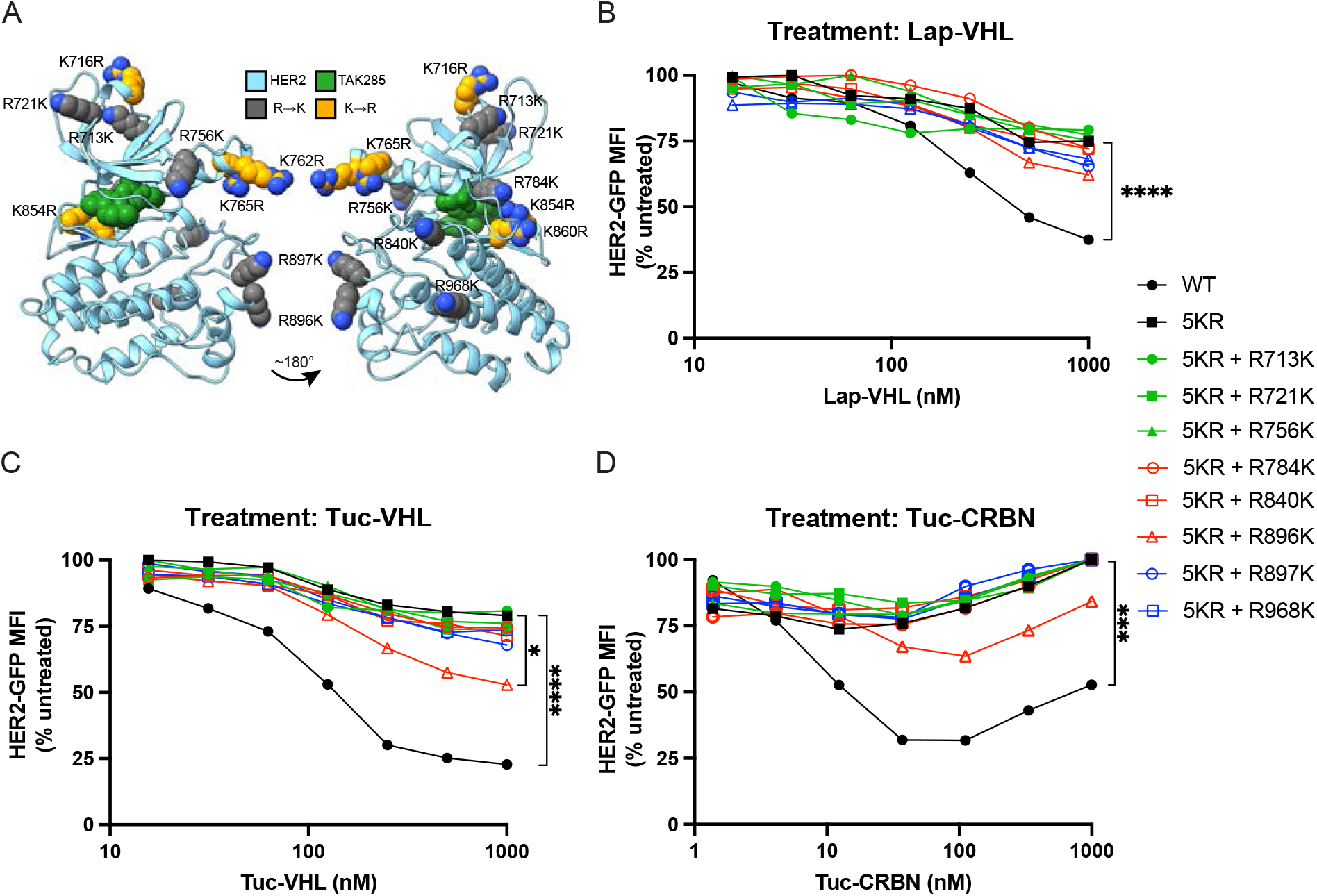
Engineered ubiquitination sites do not impact levels of degradation-resistant Her2. A. Mapping of 5KR mutations and arg-lys mutations onto the crystal structure of Her2 KD bound to TAK285. B-D. Quantification of mean Her2-GFP levels in cells treated for 20 hrs with the indicated degraders. Each data point represents a single measurement, and each panel is representative of two biological replicates. ****p < 0.0001, *** p < 0.001, *p < 0.05 calculated using two-way ANOVA corrected with Dunnett’s multiple comparisons test.

## Discussion

Structural and functional analyses of reconstituted complexes have generated important insights into how CRLs ubiquitinate their substrates, but the dynamic nature of ubiquitin poses significant challenges to detecting ubiquitinated forms in intact cells. To overcome these limitations, we used heterobifunctional degraders to induce ubiquitination of model CRL substrates and K-ε-GG enrichment and analysis to identify ubiquitination sites. By performing this analysis in a cellular system, we were able to use genetic approaches to investigate the relationship between site-specific ubiquitination and degradative outcomes.

Based on the crystal structure of lapatinib-bound EGFR, we predict that the ubiquitinated surface is oriented oppositely from the KD face that binds VHL. Consequently, a subset of lysines on this surface are likely juxtaposed with UCEs within the CRL complex. Comparing ubiquitination sites between Her2 and EGFR reveals that UCEs target lysines based on shared spatial orientation rather than strictly sequence homology. Interestingly, we also detected several constitutively ubiquitinated sites in Her2 that were unaffected by degrader treatment, aligning with findings from previous Her2 proteomics studies^22^. Our data show that degraders largely trigger additional ubiquitination of a subset of these constitutively ubiquitinated sites, suggesting that they are accessible and possess the necessary spatial orientations. This spatial requirement helps explain why degrader-dependent formation of more stable ternary complexes does not always correlate with degradation efficiency^23^, as ubiquitin transfer also depends on how the substrate interfaces with the UCE. Supporting this idea, we found that similar sites were ubiquitinated regardless of the KD binder or CRL adaptor. Intriguingly, active degraders reduced ubiquitination of K753, which is known to participate in ATP binding. Since the molecules bind the KD through the ATP binding pocket, this observation raises the possibility that K753 ubiquitination may also negatively regulate ATP binding and kinase activity. Ultimately, further structural studies of the KD-UCE interface are needed to determine how contacts between these proteins directs ubiquitin transfer.

A recent study used similar approaches to identify ubiquitination sites on the BD2 domain of BRD4, finding that multiple Lys-Arg mutations were required to reduce BD2 degradation^5^. However, because these mutations also decreased degrader potency, it is possible that they interfered with degrader binding to BD2. In contrast, our results demonstrate that lysine mutagenesis specifically affected maximum Her2 degradation (D_max_) without impairing target engagement (DC_50_), a finding consistent with the comparable inhibition of wild-type and mutant Her2 phosphorylation. Compared with VHL-dependent molecules, the magnitude of degradation by Tuc-CRBN was better correlated with the number of available ubiquitination sites, suggesting Cul4-bound UCEs ubiquitinate a more limited range of lysines.

We previously showed that degrader-dependent ubiquitination triggers RTK degradation through the endolysosomal pathway^12^. An earlier study using epidermal growth factor (EGF) to drive EGFR internalization and degradation found that multiple polyubiquitinated sites in EGFR had to be mutated to attenuate this process^24^. Intriguingly, the authors demonstrated that while ubiquitination was dispensable for EGFR internalization, it was required for subsequent trafficking of EGFR to lysosomes. In contrast, degrader-induced polyubiquitination likely regulates both initial RTK internalization and trafficking. Thus, the ability to induce ubiquitination of diverse substrates and control their ubiquitination status in time using degraders presents opportunities to study the role of ubiquitin signaling in distinct protein degradation pathways.

## Author contributions

N.O. and S.G. performed the mass spectrometry experiments and analyzed the resulting data. K.G. and M.E.F. designed and synthesized the degrader molecules. S.V. performed immunopurification assays. A.H.N. performed protein modeling. F.M. supervised the mass spectrometry efforts. D.S. supervised the project and collaborations. N.O., A.H.N, S.V. and K.B. contributed figures and wrote the manuscript. K.B. conceived the project and conducted the remaining experiments.

## Data availability

The raw mass spectrometry proteomics data have been deposited to the ProteomeXchange Consortium via the PRIDE^25^ partner repository with the dataset identifier PXD076283.

## Acknowledgements and disclosures

We thank Dr. Ian Foe and Dr. Robert Cohen (Calico Life Sciences LLC) for critical reading of the manuscript. We thank Dr. Isabelle Chiu (C4 Therapeutics, Inc.) for supporting collaborations enabling small molecule development. Calico Life Sciences LLC provided funding for all resources used in this work. The authors declare no competing interests.

## Methods

### Cell lines and culture conditions

SK-BR-3 cells, obtained from ATCC, were cultured at 37°C with 5% CO_2_ in RPMI 1640 medium containing 10% fetal bovine serum (FBS) and Glutamax (Gibco, cat. #35-050-061). SK-BR-3 cells expressing inducible wild type or mutant Her2-GFP were generated by infecting parental cells with a lentivirus encoding Her2-GFP under control of a doxycycline-inducible promoter. Virus-containing media from transfected HEK293T was added at a 1:1 v/v ratio to cells with 10 μg/ml Polybrene infection reagent. The media was changed after 24 hr, and after an additional 24 hr incubation, infected cells were selected with 2 μg/mL puromycin for 72 hr and maintained in 2 μg/mL puromycin during subsequent passages. Her2-GFP-positive cells were further enriched by fluorescence activated cell sorting using a SH800S cell sorter (Sony Biotechnology) following induction of Her2-GFP expression with 500 ng/ml doxycycline for 48 hrs.

### Plasmids

Dox-inducible Her2-GFP was generated by fusing the sequence of human Her2 to the N-terminus of GFP through a glycine-serine linker and inserting the resulting sequence between the NheI and NotI sites of pHSUSHV2, a custom dox-inducible vector encoding a puromycin-resistance cassette. Her2-GFP Lys-Arg and Arg-Lys mutants were generated using site directed mutagenesis of Her2-GFP.

### Western blotting

To prepare lysates, cells were scraped into media, pelleted by centrifugation and washed with PBS. Lysates were prepared by resuspending cell pellets in 1% SDS/water. The lysates were homogenized by shaking in a Tissuelyser II (QIAGEN) for 4 min at 30 Hz, and heated at 100°C for 5 min. Following quantification of protein concentration using Bicinchoninic acid (BCA) assay (Thermo Fisher Scientific, cat. #23225), 40 μg of lysate was combined with Laemmli buffer containing 2.5% v/v β-Mercaptoethanol (BME) and heated at 65°C for 10 min. -PAGE on a 4-20% Criterion TGX gel (Bio-Rad) at 150V for 75 min.

Proteins were transferred to 0.22 μm nitrocellulose membranes (Bio-Rad) using the Trans-Blot Turbo transfer system (Bio-Rad). To block, membranes were incubated in 5% BSA/TBST for 30 min at room temperature. Membranes were then incubated in primary antibodies overnight at 4°C, washed three times with TBST, and blotted with secondary antibodies for 30 min at room temperature. After three additional TBST washes, imaging was performed using an Odyssey M Imaging System (LI-COR).

The following antibodies were used for immunoblotting: anti-Her2 Rb mAb Clone D8F12 (Cell Signaling Technology, cat. #S920S), anti-Her2 Ms mAb Clone 44E7 (Cell Signaling Technology, cat. #2248S), anti-Phospho-Her2 (Tyr1196) mRb Clone D66B7 (Cell Signaling Technology, cat. #D66B7), anti-Ubiquitin P37 (Cell Signaling Technology, cat. #58395S), anti-Ubiquitin K48-linkage Specific Polyubiquitin Rb mAb Clone D9D5 (Cell Signaling Technology, cat. #8081S), anti-GFP Ms mAb Clone 1E10H7 (Proteintech, cat. #66002-1-Ig), anti-GAPDH (Cell Signaling Technology cat. #97166), anti-mouse IRDye 680RD (Li-COR), anti-rabbit IRDye 800 CW (Li-COR), anti-rabbit HRP-linked (Cell Signaling Technologies, cat. #7074P2) and Peroxidase AffiniPure anti-mouse (Jackson ImmunoResearch cat. #115035003). Anti-GAPDH (Cell Signaling Technologies, cat. #97166), anti-rabbit/mouse IRDye, and HRP-linked anti-rabbit/mouse antibodies were diluted 1:5000 v/v in 5% BSA/TBST. All other primary antibodies were diluted 1:1000 v/v in 5% BSA/TBST.

### Immunopurification

For each treatment condition and Her2-GFP-expressing cell line, ten 15-cm plates were treated for 48 hr with 500 ng/mL doxycycline. After a 2 hr treatment with degraders, cells were scraped into culture media, pelleted at 4°C and washed twice with cold PBS. The cell pellets were lysed in cold RIPA buffer containing 50 mM Tris-HCl pH 7.5, 150 mM NaCl, 1 mM EDTA, 0.1% SDS, 1% Triton X-100 and 1% Sodium deoxycholate, supplemented with cOmplete EDTA-free Protease Inhibitor Cocktail (Sigma, cat. #11873580001) and 5 mg/mL N-ethylmaleimide (NEM, Sigma). The lysates were rotated for 30 min at 4°C and cleared by centrifugation for 15 min at 20 k x g. BCA assay was used to quantify the protein concentration of the supernatant. To perform the immunopurification, 25 μL of Chromotek GFP-Trap agarose bead slurry (Proteintech Group, cat. #gta) was washed with twice with RIPA buffer and added to 15 mg of lysate per sample. The combined samples were rotated at 4°C for 2 hr. Beads were pelleted at 2000 x g, washed four times with cold RIPA buffer and twice with cold wash buffer containing 50 mM Tris-HCl pH 7.5, 150 mM NaCl, and 0.5 mM EDTA. After removal of supernatant, 2x Laemmli buffer was added to the beads, and bound proteins were eluted by boiling for 5 min. Eluted proteins were subsequently combined with 2.5% BME, heated at 65°C for 10 min, and separated as described above. Proteins were transferred to 0.45 μm PVDF membrane (Bio-Rad), blotted with the indicated primary antibodies at 4°C ON, and visualized by blotting with HRP-linked anti-Rabbit or anti-Mouse secondary antibodies for 30 min at RT, followed by a 5 min incubation in SuperSignal West Pico PLUS chemiluminescent substrate (Thermo Fisher Scientific, cat. #34580).

### Flow cytometry

The LSRFortessa X-20 cell analyzer (BD Biosciences) was used to perform all flow cytometric analyses. For analysis of Her2-GFP levels, cells were seeded into 96-well tissue culture plates and treated with 500 ng/mL doxycycline for 48 hr. After a 20 hr treatment with degraders, the cell culture media was aspirated, and all wells were washed with PBS. Cells were trypsinized with 100 uL TrypLE at 37°C for 3 min and transferred to a V-bottom plate containing 100 uL of complete media per well. The cells were pelleted at 500 x g for 3 min, washed 1X with PBS, and fixed in 100 uL of 0.5% paraformaldehyde/PBS.

FlowJo analysis software (FlowJo, LLC, v.10.10.0) was used to select the GFP-positive population and calculate their geometric mean fluorescence intensity (MFI). These values were normalized by subtracting the MFI of the GFP-negative population, and the percent Her2-GFP remaining was calculated by dividing by the normalized MFI of the untreated population. GraphPad PRISM (Dotmatics, v10.6.1) was used to plot values and calculate statistical significance.

### Mass spectrometry

For each replicate, fifteen 15-cm plates of SK-BR-3 cells were treated with degraders for 3 hr, washed twice with cold PBS at 4°C, and scraped into cold PBS. Cells were pelleted by centrifugation at 500 x g for 5 min, prior to freezing at -80°C.

Analysis of diGly-enriched samples was performed across two studies (see supplementary methods). The Lap-VHL study comprised a 6-plex TMT study with two replicates each of Lap-VHL-treated cells, Lap-VHL_Inactive_-treated cells and DMSO-treated cells. The Tuc-VHL/Tuc-CRBN study comprised a 16-plex TMT study with three replicates each of Tuc-VHL, Tuc-VHL_Inactive_, Tuc-CRBN and Tuc-CRBN_Inactive_-treated cells, and four replicates of DMSO-treated cells. In the Lap-VHL study, the acquisition of 101,999 MS2 spectra yielded 25,938 unique peptides. After the removal of contaminants and decoy entries (1% FDR), 25,191 unique peptides remained, of which 14,439 were identified as unique di-glycine (K-GG) modified peptides. In the Tuc-VHL/Tuc-RBN study, 172,600 MS2 spectra were acquired, resulting in 11,676 unique peptides. Following the exclusion of contaminants and decoys, 11,310 unique peptides were identified, including 6,762 unique di-glycine modified peptides. In this context, unique peptides refer to distinct sequences and modification states (e.g., a sequence with and without methionine oxidation is counted as two distinct peptides).

### Structural analysis

Molecular graphics and analyses were performed with UCSF ChimeraX version 1.9, developed by the Resource for Biocomputing, Visualization, and Informatics at the University of California, San Francisco, with support from National Institutes of Health R01-GM129325 and the Office of Cyber Infrastructure and Computational Biology, National Institute of Allergy and Infectious Diseases^26^.

AlphaFold2 (AF2) models of Her2 (AF-P04626-F1-model_v6.pdb; residues Q709-V988) and EGFR (AF-P00533-F1-model_v6.pdb; residues Q701–V980) were downloaded from UniProt (Consortium, 2024) and used to fill in missing residues. Mutations depicted in Figure 4 were introduced computationally in UCSF ChimeraX using the Rotamers tool with the Dunbrack rotamer library, and the most probable rotamer was selected for visualizations^27^. These side-chain conformations were used for illustrative purposes only and were not derived from structural or energetic calculations.

To depict TAK-285 or lapatinib bound within the Her2 or EGFR kinase domains, published X-ray crystal structures were used as ligand-bound references: TAK-285:Her2 from PDB ID no. 3RCD^17^ and lapatinib:EGFR from PDB ID no. 1XKK^18^. The ligand-bound X-ray structures were superimposed onto the corresponding AF2 models using the MatchMaker utility in UCSF ChimeraX. The crystallographic protein model was then hidden to allow visualization of the small molecule within the context of the full kinase-domain representation.

Small-molecule and heterobifunctional degrader structures were drawn in ChemDraw version 22.2.0.3348 (Revvity signals) and imported into Adobe Illustrator (v30.2.1) for figure assembly.

## Supplemental Information

Document S1. Figures S1-S3, supplemental chemical synthesis and mass spectrometry methods and chemical analysis. Data S1. Mass spectrometry IDs and quantification.

