## Supplemental Figures and Methods for "Site-specific combinatorial ubiquitination drives the targeted degradation of plasma membrane proteins"

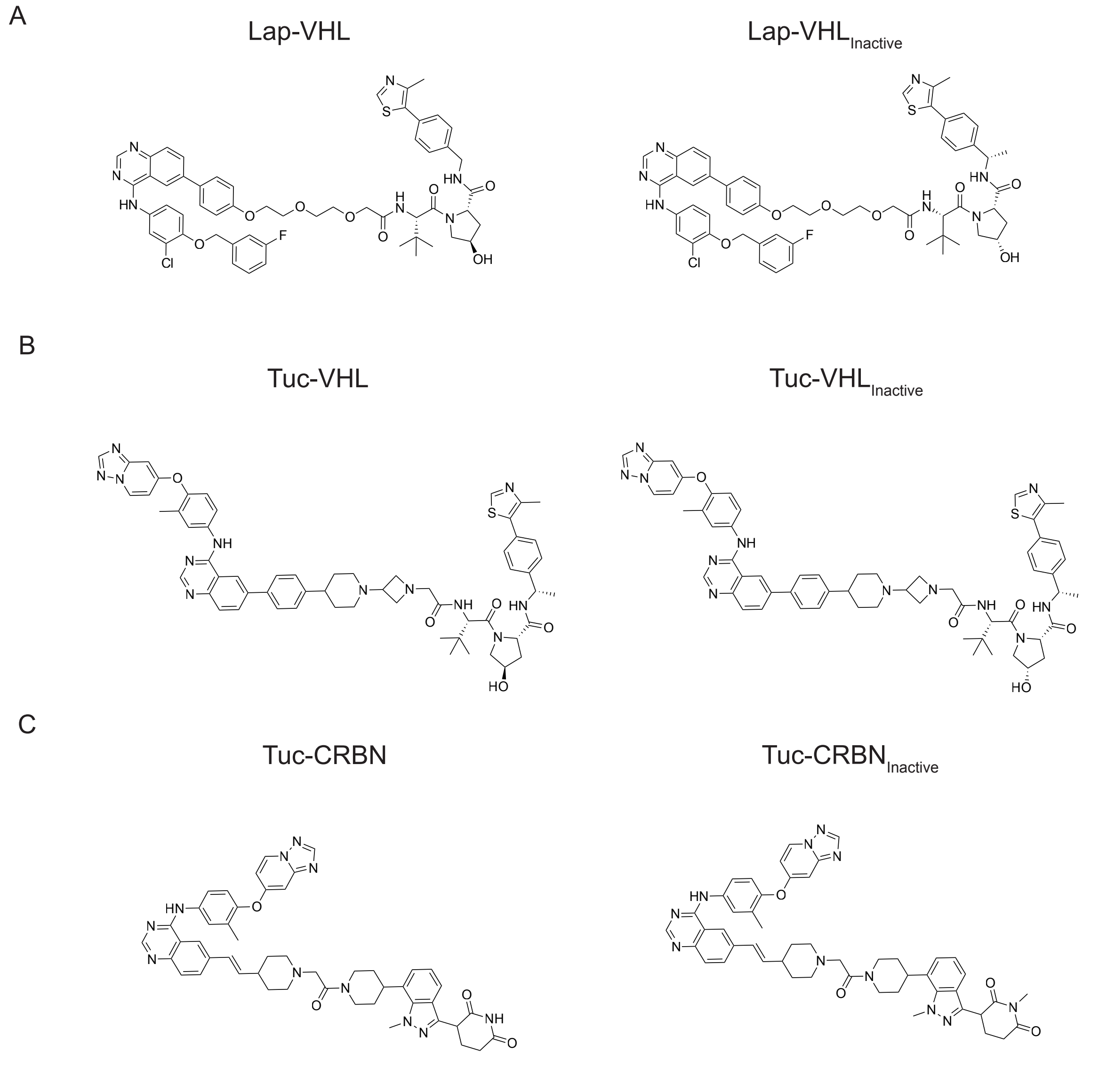


**Figure S1. Structures of heterobifunctional degraders used in this study, related to Figure 1.**

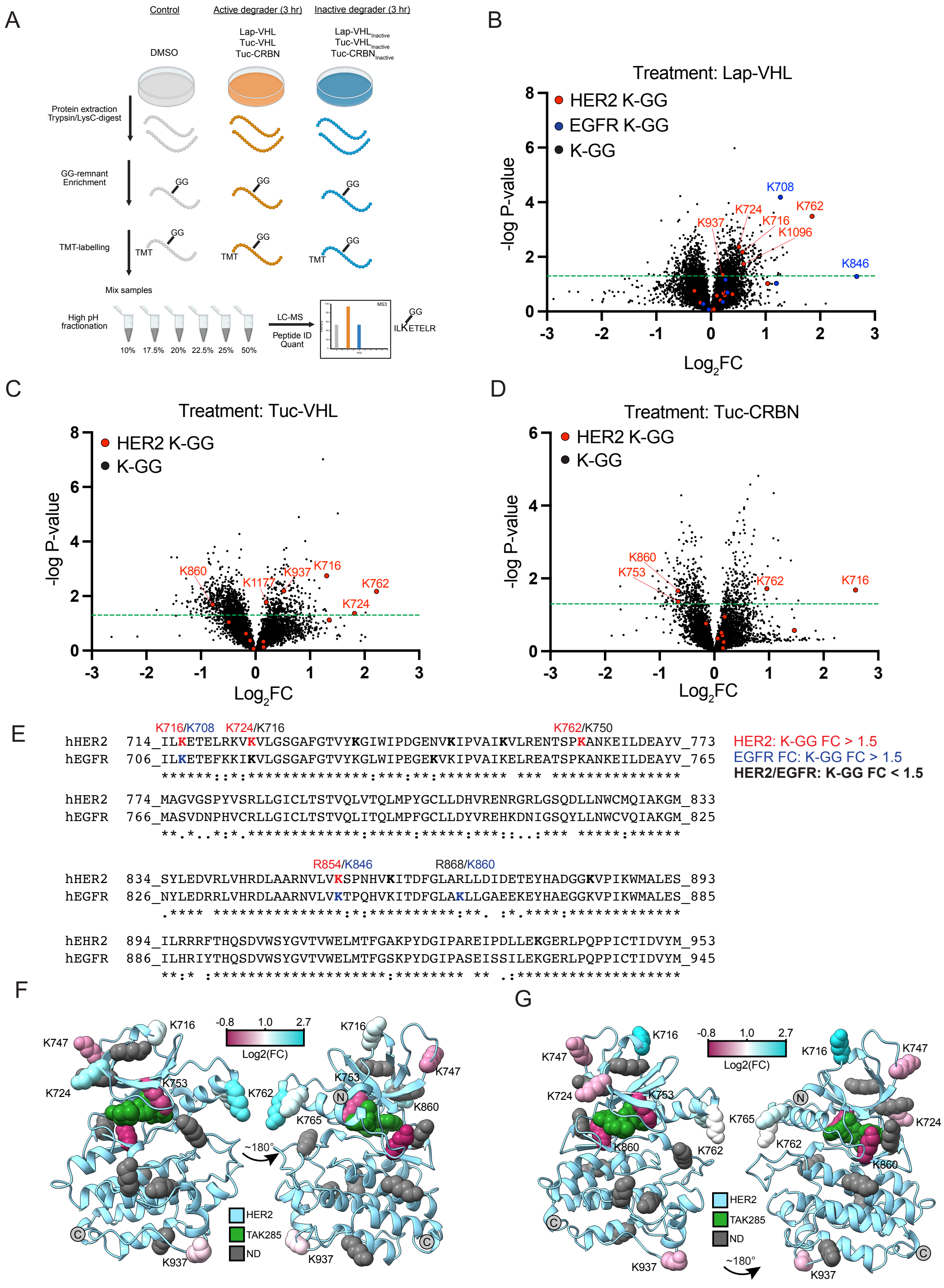
A
Control
Active degrader (3 hr)
Inactive degrader (3 hr)
DMSO
Lap-VHL
Tuc-VHL
Tuc-CRBN
Protein extraction
Trypsin/LysC-digest
GG-remnant Enrichment
TMT-labelling
Mix samples
High pH fractionation
10% 17.5% 20% 22.5% 25% 50%
LC-MS
Peptide ID Quant
ILKETELR
B
Treatment: Lap-VHL
HER2 K-GG
EGFR K-GG
K-GG
-log P-value
Log2FC
K708 K762 K724 K716 K937 K1096 K846
C
Treatment: Tuc-VHL
HER2 K-GG
K-GG
K860 K1177 K937 K716 K762 K724
D
Treatment: Tuc-CRBN
K860 K753 K762 K716
E
K716/K708 K724/K716 K762/K750
hHER2 714_ILKETELRKVKVLGSGAFGTVYKGIWIPDGENVKIPVAIKVLRENTSPKANKEILDEAYV_773
hEGFR 706_ILKETEFKKIKVLGSGAFGTVYKGLWIPEGEKVKIPVAIKELREATSPKANKEILDEAYV_765
HER2: K-GG FC > 1.5
EGFR FC: K-GG FC > 1.5
HER2/EGFR: K-GG FC < 1.5
hHER2 774_MAGVGSPYVSRLLGICLTSTVQLVTQLMPYGCLLDHVRENRGRLGSQDLLNWCMQIAKGM_833
hEGFR 766_MASVDNPHVCRLLGICLTSTVQLITQLMPFGCLLDYVREHKDNIGSQYLLNWCVQIAKGM_825
R854/K846 R868/K860
hHER2 834_SYLEDVRLVHRDLAARNVLVKSPNHVKITDFGLARLLDIDETEYHADGGKVPIKWMALES_893
hEGFR 826_NYLEDRRLVHRDLAARNVLVKTPQHVKITDFGLAKLLGAEEKEYHAEGGKVPIKWMALES_885
hEHR2 894_ILRRRFTHQSDVWSYGVTVWELMTFGAKPYDGIPAREIPDLLEKGERLPQPPICTIDVYM_953
hEGFR 886_ILHRIYTHQSDVWSYGVTVWELMTFGSKPYDGIPASEISSILEKGERLPQPPICTIDVYM_945
F
-0.8 1.0 2.7
Log2(FC)
K716 K747 K724 K753 K762 K765 K860 K937
~180°
HER2
TAK285
ND
G
K716 K747 K724 K753 K765 K762 K860 K937
~180°
HER2
TAK285
ND

**Figure S2. Identification of ubiquitination sites using diglycine remnant motif proteomics, related to Figure 2.** A. Schematic showing the experimental workflow to profile ubiquitination sites using di-glycine remnant motif (K-ε-GG) proteomics. Protein lysates from degraders-treated cells were digested with trypsin, and peptides containing lysines conjugated to the ubiquitin GG remnant motif (K-GG) were immunopurified using anti-K-GG antibodies. Peptides from different treatments were multiplexed using TMT labeling and abundance ratios were determined using mass spectrometric analysis. B-D. Fold change in K-GG site abundance in active degrader-treated cells. Samples from the corresponding inactive degrader-treated cells were grouped with DMSO-treated samples to calculate fold change and perform statistical analysis. B was run as a 6-plex TMT experiment containing two replicates for each treatment group. C,D were run as a combined 16-plex TMT experiment containing three replicates for each degrader treatment and four replicates for DMSO. P-values were determined using a two-tailed Welch's t-test on the normalized data. E. Sequence alignment between Her2 and EGFR kinase domains showing ubiquitination sites that exhibit the largest fold changes. F,G. Mapping of identified K-GG sites from Tuc-VHL (F) or Tuc-CRBN (G) treated cells onto the crystal structure of Her2 bound to TAK285.

Figure S3

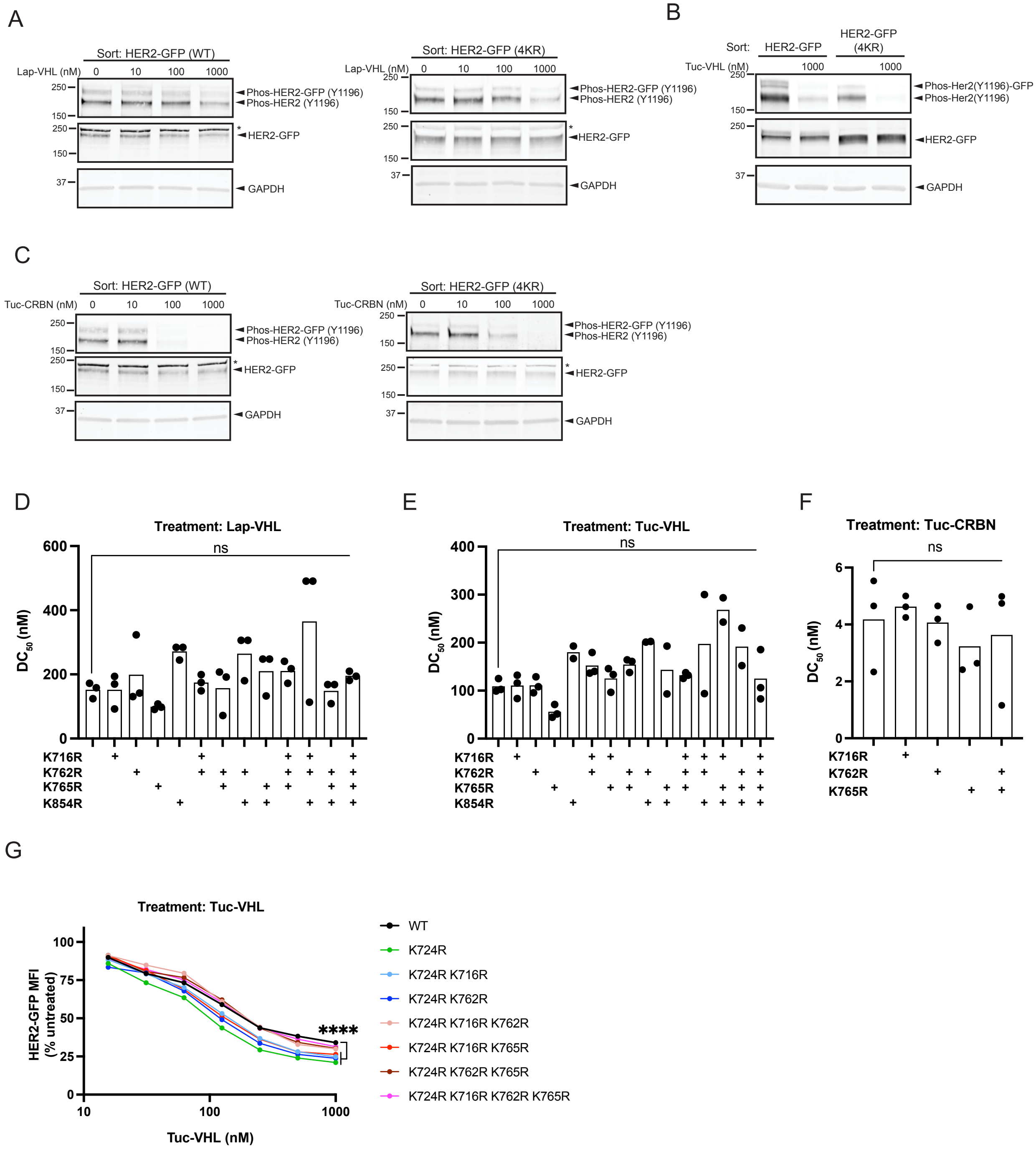

A
Sort: HER2-GFP (WT)
Lap-VHL (nM) 0 10 100 1000
Phos-HER2-GFP (Y1196)
Phos-HER2 (Y1196)
HER2-GFP
GAPDH
Sort: HER2-GFP (4KR)
B
Sort: HER2-GFP HER2-GFP (4KR)
Tuc-VHL (nM) 1000 1000
Phos-Her2(Y1196)-GFP
Phos-Her2(Y1196)
C
Tuc-CRBN (nM) 0 10 100 1000
D
Treatment: Lap-VHL
DC50 (nM)
ns
K716R
K762R
K765R
K854R
E
Treatment: Tuc-VHL
F
Treatment: Tuc-CRBN
G
Treatment: Tuc-VHL
HER2-GFP MFI (% untreated)
Tuc-VHL (nM)
****
WT
K724R
K724R K716R
K724R K762R
K724R K716R K762R
K724R K716R K765R
K724R K762R K765R
K724R K716R K762R K765R

**Figure S3. Characterization of Her2-GFP containing lysine to arginine mutations, related to figure 3.** A-C. Analysis of Her2-GFP and Her2 (4KR)-GFP phosphorylation status in lysates from cells treated for 2 hrs with the indicated molecules. After treatment, GFP-positive cell populations were sorted and phosphorylation of endogenous Her2 and Her2-GFP was determined using western blot. An anti-GFP antibody was used to detect total HER2-GFP. The blots show the results of single experiments. *, non-specific band. D-F. The concentration of degraders required for half maximal degradation ($DC_{50}$) was determined from measurements in Figures 3G-K. Each data point represents a biological replicate. ns, $p > 0.05$. G. Flow cytometric analysis of wild type and mutant Her2-GFP levels in cells treated for 20 hr with 1 μM Tuc-VHL. ****$p < 0.0001$.

**Supplemental methods**

Mass spectrometry

*Cell Lysis and Protein Extraction*

Cells were resuspended in lysis buffer (8 M Urea, 150 mM NaCl, 50 mM HEPES pH 7.2) supplemented with protease inhibitors (cOmplete ULTRA, Roche) and 10 mM N-ethylmaleimide (NEM) as a deubiquitinating enzyme (DUB) inhibitor and Lysis was achieved via 20 passages through a 21-gauge syringe followed by sonication (5 cycles of 10 s on/10 s off). Protein concentrations were determined via BCA Assay Kit (Pierce). Lysates were reduced with 10 mM TCEP for 1.5 h at room temperature and alkylated with 15 mM NEM for 30 min. NEM was utilized specifically to avoid double-alkylation artifacts associated with iodoacetamide that mimic the di-glycine remnant mass shift. Excess NEM was quenched with 10 mM DTT for 15 min. Proteins were recovered by methanol/chloroform precipitation. The resulting pellets were resuspended in 8 M Urea/50 mM HEPES (pH 8.5) and subsequently diluted to 1 M Urea for proteolysis. Samples were digested with LysC (1:250) for 16 h at room temperature, followed by Trypsin/LysC (final total protease ratio 1:100) for 6 h at 37°C. Digestion was quenched with 0.5% TFA, and peptides were desalted using C18 solid-phase extraction (Sep-Pak, Waters).

*Immunoaffinity Purification (IAP) of Di-Gly Peptides*

Enrichment of ubiquitinated peptides was performed using the PTMScan HS Ubiquitin Remnant Motif (K-epsilon-GG) Kit (Cell Signaling Technology). For each sample, 4 mg of dried peptides were resuspended in 1.5 mL IAP-HS bind buffer. The solution was cleared by centrifugation (10,000xg, 5 min), and the supernatant was incubated with 20 µL of pre-washed antibody-bead slurry for 2 h at 4°C on an end-over-end rotator. Beads were washed four times with HS IAP Wash Buffer and twice with LC-MS grade water. Enriched peptides were eluted twice with 50 µL of 0.2% TFA, desalted via C18 StageTips[1], and then were dried completely (SpeedVac).

*TMTpro Labeling*

Enriched peptides were resuspended in 18 µL of 200 mM EPPS (pH 8.5) and 4 µL anhydrous ACN. Labeling was performed by adding 7.2 µg of TMTpro 16-plex reagents for 1 h at room temperature. Reactions were quenched with 5 µL of 5% hydroxylamine for 15 min and dried.

*High-pH Reversed-Phase (HPRP) Fractionation*

Labeled peptides were fractionated using a High pH Reversed-Phase Peptide Fractionation Kit (Thermo Fisher Scientific) according to the manufacturer's instructions with some minor modifications. For both experiments, columns were equilibrated with ACN and 0.1% TFA. Samples were loaded in 0.1% TFA and washed with water. Elution was performed using varying concentrations of ACN in 0.1% triethylamine. For the Lap-VHL study, step elutions of 17.5%, 20%, 22.5%, 25%, 30%, and 70% ACN were collected. For the Tuc-VHL/Tuc-CRBN study, elutions of 5%, 10%, 17.5%, 20%, 22.5%, 25%, and 50% ACN were collected; the 5% and 50% fractions were subsequently merged to yield six final fractions for both experiments. All fractions were dried, desalted using homemade StageTips[1], dried completely (SpeedVac) and finally resuspended in 5% ACN/5% formic acid for LC-MS/MS analysis.

*LC-MS/MS Data Acquisition*

Peptides were analyzed by LC-MS/MS using either an Orbitrap Fusion Lumos (HDK experiment) coupled to an EASY-nLC 1200 ultra-high-pressure liquid chromatography (UPLC) system or an Orbitrap Eclipse (HEO experiment) mass spectrometer equipped with a FAIMS Pro (Field Asymmetric Ion Mobility Spectrometry) interface and coupled to an UltiMate 3000 HPLC system (Thermo Fisher Scientific). In both experiments, peptides were separated on an Aurora Series emitter column (25 cm × 75 µm i.d., C18; IonOpticks) at a constant flow rate of 300 nL/min. For the Lumos runs, the gradient transitioned from 8% to 28% buffer B (95% ACN, 0.1% formic acid) over 165 min, followed by an increase to 95% B over 7 min. For the Eclipse , the 185-minute method employed an initial equilibration at 1%

buffer B followed by a gradient increase to 6% B at 10 min. Subsequently, a linear increase reached 18% B at 145 min , 30% B at 158 min , and 35% B at 162.5 min. The run concluded with a high-organic wash at 90% B for 8 min before returning to initial conditions.

The scan sequence for both instruments utilized a Top-speed data-dependent acquisition (DDA) method[2] with some adjustments. In brief, the Lumos was operated with a 5 s cycle time, while the Eclipse utilized a 1.25 s cycle time per FAIMS compensation voltage (CV). On the Eclipse, four CVs were interrogated sequentially: -40 V, -50 V, -60 V, and -70 V. The sequence began with an FTMS1 scan in the Orbitrap at a resolution of 120,000. The mass range was set to 400–1250 m/z for the Lumos and 400–1600 m/z for the Eclipse , with absolute AGC targets of $5 \times 10^5$ and $4.0 \times 10^5$, respectively. Maximum injection times were set to 100 ms for the Lumos or utilized Auto mode on the Eclipse , with an RF lens of 30%. Precursors were filtered by charge state (3–7 for Lumos; 3–6 for Eclipse) , and monoisotopic peak assignment was enforced using MIPS mode. Interrogated precursors were excluded using a dynamic window of 60 s duration with a mass tolerance of ± 7.5 ppm (Lumos) or ± 10 ppm (Eclipse).

For MS2 analysis, precursors were isolated using the quadrupole with an isolation window of 0.5 Th (Lumos) or 0.7 Th (Eclipse). On the Lumos, precursors were fragmented by collision-induced dissociation (CID) at a normalized collision energy (NCE) of 34% (activation Q = 0.25). Spectra were analyzed in the Orbitrap at 15,000 resolution with an AGC target of $1 \times 10^5$ and a 300 ms maximum injection time. On the Eclipse, precursors were fragmented by high-energy collision-induced dissociation (HCD) at an NCE of 30%. These spectra were analyzed in the Orbitrap at 15,000 resolution with an absolute AGC target of $1.0 \times 10^5$ and a maximum injection time of 250 ms.

Following each MS2 spectrum, a synchronous-precursor-selection (SPS)-MS3 scan was collected for TMT reporter ion quantification. The top 10 most intense ions in the MS2 spectrum were selected for MS3 analysis via Multi-notch Isolation with a 2 m/z isolation window. Precursors were fragmented by HCD at an NCE of 45% and analyzed in the Orbitrap across a 100–500 m/z range with an absolute AGC target of $1.0 \times 10^5$. For the Lumos, resolution was set to 60,000 with a 500 ms maximum injection time, while the Eclipse utilized a resolution of 50,000 and a maximum injection time of 250 ms.

*Data processing*

Mass spectrometric raw data files were converted to mzXML format using the MSConvert program and processed through a compilation of in-house software to correct monoisotopic m/z measurements and erroneous peptide charge state assignments[3]. Fragment spectra were assigned using the Comet[4] algorithm (version 2023.01 rev. 0) and queried against a human target-decoy protein sequence database consisting of the *Homo sapiens* UniProt database (6/4/2020) concatenated with a decoy database composed of all protein sequences in reversed order, as well as common contaminants and their reversed decoys[5]. A precursor ion tolerance of 20 ppm was employed, and the product ion tolerance was set to 0.02 Da for fragment binning. Peptides were required to possess full enzyme specificity (LysC/Trypsin) at both N- and C-termini, with up to 4 missed cleavages allowed. Static modifications included NEM-alkylation on cysteines (+125.04767 Da) and TMTpro labels on both peptide N-termini and lysine residues (+304.207146 Da). Variable modifications were set for methionine oxidation (+15.99491 Da) and lysine ubiquitination (di-glycine remnant; +114.04293 Da), with a maximum of 4 variable modifications permitted per peptide. Peptides shorter than seven amino acid residues in length were discarded. Following the database search, peptide-spectrum matches were filtered to a 1% peptide-level FDR. Known false positives (i.e. decoys) and contaminants were removed. The confidence of the site localizations was determined by ModScore, which is similar to Ascore. It employs a probabilistic algorithm that considers the possible modification (di-glycine remnant) sites in a modified peptide and uses the presence of experimental fragment ions unique to each site to find the best match[6]. Sites with ModScore>13 ($p < 0.05$) were considered as confidently localized. To facilitate quantitative comparison across the high-dimensional ubiquitination datasets, site-level signal-to-noise (S/N) ratios were subjected to global median-scaling normalization across all TMT channels. Statistical significance was evaluated using a two-tailed Welch's t-test on the normalized data and either a fold-change or nominal significance threshold of $p < 0.05$ was employed for targeted analysis of the ERBB family (ERBB2, ERBB3 and EGFR).

Chemical synthesis

Lap-VHL was purchased from BioTechne (Catalog #: 7262) and used as received.

Synthetic procedures for Lap-$VHL_{Inactive}$, Tuc-VHL, and Tuc-$VHL_{Inactive}$ have been previously reported.[7]

*Synthesis of Intermediate 7*

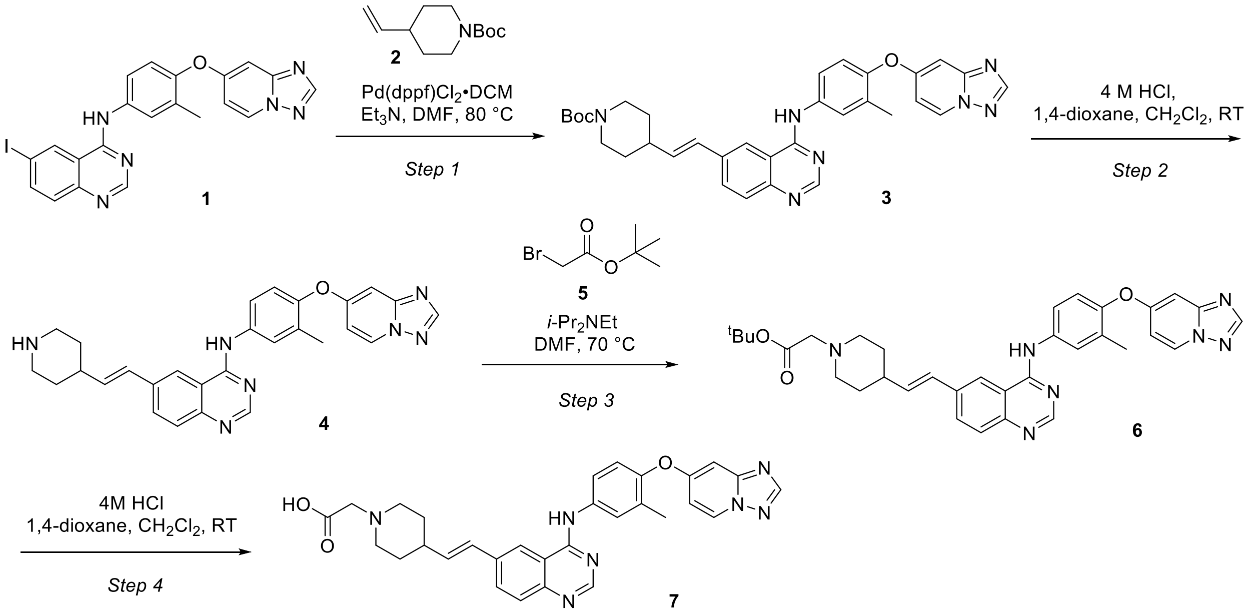


Step 1: To a room temperature solution of 6-iodo-*N*-[3-methyl-4-([1,2,4] triazolo[1,5-a] pyridin-7-yloxy) phenyl]quinazolin-4-amine[1] 1 (1.0 g, 2.02 mmol) and *tert*-butyl 4-vinylpiperidine-1-carboxylate 2 (513 mg, 2.43 mmol) in anhydrous DMF (25 mL) was added triethylamine (4.09 g, 40.46 mmol, 5.64 mL) under nitrogen atmosphere. The reaction mixture was purged with nitrogen gas for 10 min, then Pd(dppf)$Cl_2 \cdot CH_2Cl_2$ (165 mg, 202.31 µmol) was added at room temperature. The resulting reaction mixture was heated to 80 °C and stirred for 16 h. The reaction mixture was diluted with water (25 mL) and extracted with ethyl acetate (3 x 30 mL). The combined organic layer was dried over anhydrous sodium sulfate, filtered, and concentrated under reduced pressure to afford the crude compound, which was purified by silica gel chromatography (0-15% methanol in dichloromethane) to afford *tert*-butyl 4-[(*E*)-2-[4-[3-methyl-4-([1,2,4] triazolo[1,5-a] pyridin-7-yloxy)anilino]quinazolin-6-yl]vinyl]piperidine-1-carboxylate 3 (1.0 g, 1.14 mmol, 56% yield, 65.7% purity) as a yellow solid. LC-MS (ESI) *m*/*z* calculated for $C_{33}H_{35}N_7O_3$ $[M + H]^+$ 578.28, found: 578.2.

Step 2: To a stirred solution of 3 (1.0 g, 1.73 mmol, 1.0 eq) in dichloromethane (8.0 mL) was added 4M HCl in 1,4-dioxane (6.49 mL, 25.97 mmol, 1.0 eq) at room temperature under nitrogen atmosphere. The reaction mixture was stirred at room temperature for 3 h. The reaction mixture was concentrated under reduced pressure, and the resultant crude product was triturated with methyl *tert*-butyl ether (10 mL) and dried in vacuo to afford (*E*)-*N*-(4-([1,2,4]triazolo[1,5-a]pyridin-7-yloxy)-3-methylphenyl)-6-(2-(piperidin-4-yl)vinyl)quinazolin-4-amine 4 (1.0 g, 1.54 mmol, 88.90% yield, 79.1% purity, HCl salt) as a yellow solid. LC-MS (ESI) *m*/*z* calculated for $C_{28}H_{27}N_7O$ $[M + H]^+$ 478.23, found: 478.2.

Step 3: To a solution of 4 (890 mg, 1.67 mmol, HCl salt) in DMF (10 mL) were added *N*,*N*-diisopropylethylamine (1.46 mL, 8.36 mmol, 1.46 mL) followed by tert-butyl 2-bromoacetate 5 (326.31 mg, 1.67 mmol, 245 µL) at room temperature under nitrogen atmosphere. The reaction mixture was stirred at 70 °C for 16 h, then diluted with ethyl acetate (100 mL) and washed with water (2 x 100 mL). The organic layer was separated, dried over anhydrous sodium sulfate, filtered, and concentrated under reduced pressure to afford the crude product, which was purified by silica gel column chromatography (0-10% MeOH in dichloromethane) to afford *tert*-butyl (*E*)-2-(4-(2-(4-((4-([1,2,4]triazolo[1,5-a]pyridin-7-yloxy)-3-methylphenyl)amino)quinazolin-6-yl)vinyl)piperidin-1-yl)acetate 6 (310 mg, 366.74 µmol, 62.84% yield, 70% purity) as a brown viscous liquid. LC-MS (ESI) *m/z* calculated for $C_{34}H_{37}N_7O_3$ $[M + H]^+$ 592.3, found: 592.2.

Step 4: To a 0 °C solution of 6 (310 mg, 523.9 µmol, 1.0 eq.) in dichloromethane (3.5 mL) was added 4M HCl in 1,4-dioxane (2.0 mL, 523.9 µmol, 1.0 eq). The resultant reaction mixture was stirred at room temperature for 3 h. The reaction mixture was concentrated under reduced pressure, triturated with hexane/methyl *tert*-butyl ether, and dried in vacuo to afford (*E*)-2-(4-(2-(4-((4-([1,2,4]triazolo[1,5-a]pyridin-7-yloxy)-3-methylphenyl)amino)quinazolin-6-yl)vinyl)piperidin-1-yl)acetic acid 7 (250 mg, 349.62 µmol, 66.73% yield, 80% purity, HCl salt) as a brown solid. LC-MS (ESI) *m/z* calculated for $C_{30}H_{29}N_7O_3$ $[M + H]^+$ 536.24, found: 536.0.

*Synthesis of Intermediate 12*

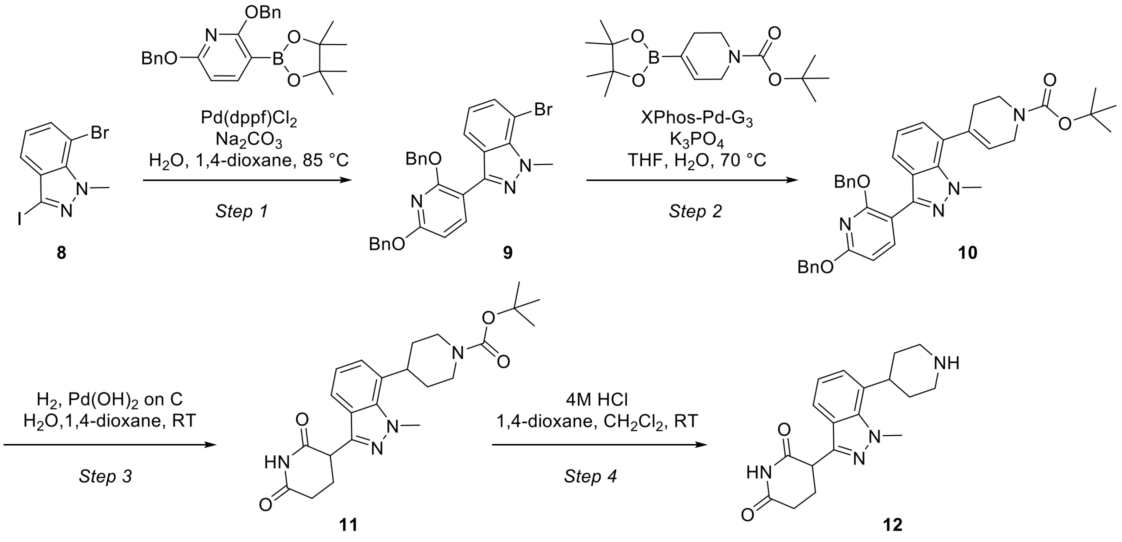


Step 1: To solution of 7-bromo-3-iodo-1-methyl-indazole 8 (6 g, 13.55 mmol, 1.0 eq) in water (20 mL) and 1,4-dioxane (60 mL) were added 2,6-dibenzyloxy-3-(4,4,5,5-tetramethyl-1,3,2-dioxaborolan-2-yl)pyridine (8.48 g, 20.33 mmol, 1.5 eq) and sodium carbonate (2.87 g, 27.10 mmol, 2.0 eq) at room temperature under nitrogen atmosphere. The reaction mixture was degassed by bubbling nitrogen through the solution for 10 min. Pd(dppf)$Cl_2$ (331.98 mg, 406.52 µmol, 0.03 eq) was added, and the reaction mixture was further degassed for 5 min then stirred at 85 °C for 16 h. The reaction mixture was poured into ice water. The aqueous layer was extracted with EtOAc (3 x 300 mL), and the combined organic layer was washed with brine (200 mL), dried over anhydrous sodium sulfate, and filtered. The filtrate was concentrated under reduced pressure to afford a crude residue which was purified by silica gel chromatography (0-20% ethyl acetate in n-hexane) to provide 7-bromo-3-(2,6-dibenzyloxy-3-pyridyl)-1-methyl-indazole 9 (3.2 g, 6.04 mmol, 44.6% yield, 94.5% purity) as an off white solid. LC-MS (ESI) *m/z* calculated for $C_{27}H_{22}BrN_3O_2$ $[M + H]^+$ 502.10, found 502.2.

Step 2: To a solution of 7-bromo-3-(2,6-dibenzyloxy-3-pyridyl)-1-methyl-indazole 9 (0.1 g, 199.85 µmol, 1.0 eq) and *tert*-butyl 4-(4,4,5,5-tetramethyl-1,3,2-dioxaborolan-2-yl)cyclohex-3-ene-1-carboxylate (73.92 mg, 239.82 µmol, 1.2 eq) in water (1 mL) and THF (4 mL) was added XPhos-Pd-G3 (16.92 mg, 19.98 µmol, 0.1 eq), and the resulting reaction mixture was purged with nitrogen for 10 minutes. Tripotassium phosphate (106.05 mg, 499.61 µmol, 2.5 eq) was added, and the reaction mixture was stirred at 70 °C for 12 hr. The reaction mixture was cooled to room temperature and diluted with ethyl acetate (70 mL) and water (20 mL). The aqueous layer was extracted with ethyl acetate (2 x 40 mL). The combined organic layers were washed with water and brine, dried over anhydrous $Na_2SO_4$, filtered and concentrated under reduced pressure. The crude product thus obtained was purified by silica gel chromatography (0-100% ethyl acetate in petroleum ether) to afford *tert*-butyl 4-[3-(2,6-dibenzyloxy-3-pyridyl)-1-methyl-indazol-7-yl]-3,6-dihydro-2*H*-pyridine-1-carboxylate 10 (55 mg, 82.13 µmol, 41.10% yield, 90% purity), as a brown solid.

Step 3: To a solution of *tert*-butyl 4-[3-(2,6-dibenzyloxy-3-pyridyl)-1-methyl-indazol-7-yl]-3,6-dihydro-2*H*-pyridine-1-carboxylate 10 (2.5 g, 4.15 mmol, 1.0 eq) in 1,4-dioxane (25 mL) was added palladium hydroxide on carbon, 20 wt.% wetted with 50% water (2.5 g) under inert atmosphere at room temperature. The reaction was hydrogenated with hydrogen bladder pressure for 24 h at room temperature. The reaction mixture was filtered through a pad of Celite®, and the Celite® pad was washed with 1,4-dioxane and ethyl acetate. The filtrate was concentrated under reduced pressure, and the crude product was purified by reverse phase chromatography [Column: Redisep Rf Gold ® reversed-phase C18-teledyne ISCO, 100 g; Mobile phase A: 10 mm ammonium bicarbonate in millQ-water ; Mobile phase B : Acetonitrile; Flow rate: 15 mL/min] to afford *tert*-butyl 4-[3-(2,6-dioxo-3-piperidyl)-1-methyl-indazol-7-yl]piperidine-1-carboxylate 11 (800 mg, 1.85 mmol, 44.52% yield, 98.45% purity) as a white solid. LCMS (ES-): *m*/*z* calculated for $C_{23}H_{30}N_4O_4$ $[M - H]^-$ 425.2, found 425.0.

Step 4: To a stirred solution of *tert*-butyl 4-[3-(2,6-dioxo-3-piperidyl)-1-methyl-indazol-7-yl]piperidine-1-carboxylate 11 (50 mg, 98.47 µmol, 1.0 eq) in dichloromethane (2 mL) was added 4M HCl in 1,4-dioxane (3.59 mg, 98.47 µmol, 1.0 eq) at 0 °C. The reaction mixture was stirred at 25 °C for 2 hr. The reaction mixture was concentrated under reduced pressure to afford a crude product which was washed with methyl *tert*-butyl ether (2 x 3 mL) and dried in vucuo to afford 3-[1-methyl-7-(4-piperidyl)indazol-3-yl]piperidine-2,6-dione 12 (30 mg, 80.93 µmol, 82.19% yield, 97.89% purity) as an off white solid. (325.0 $[M+H]^+$) with 97.89% purity. LCMS (ES-): *m*/*z* calculated for $C_{18}H_{21}N_4O_2$ $[M - H]^-$ 325.17, found 325.0.

*Synthesis of Tuc-CRBN*

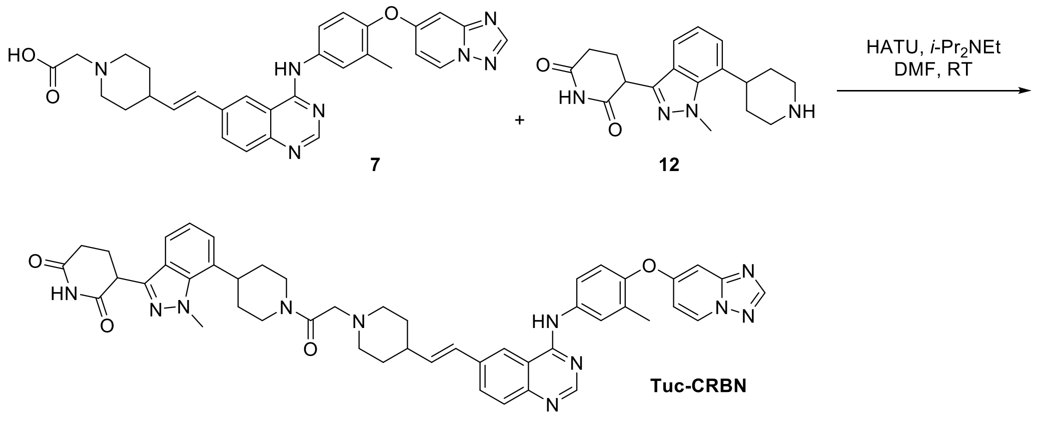

To a solution of 2-[4-[(*E*)-2-[4-[3-methyl-4-([1,2,4]triazolo[1,5-a]pyridin-7-yloxy)anilino]quin-azolin-6-yl]vinyl]-1-piperidyl]acetic acid 7 (1.4 g, 2.61 mmol 1.0 eq.) and 3-[1-methyl-7-(4-piperidyl)indazol-3-yl]piperidine-2,6-dione 12 (0.853 g, 2.35 mmol, 0.9 eq.) in anhydrous DMF (15 mL) was added *N*,*N*-diisopropylethylamine (2.28 mL, 13.07 mmol, 5.0 eq), and the resulting solution was stirred for 15 minutes at ambient temperature under nitrogen atmosphere. HATU (1.19 g, 3.14 mmol) was added and stirring was continued at ambient temperature for 2 h. The reaction mixture was quenched with ice-cold water (5 mL) and stirred for 10 minutes. The precipitated solid was filtered and dried in vacuo to afford the crude compound. The crude product was purified by reverse phase column chromatography (Column: Redisep $R_f$ Gold C18 100 g, Mobile phase A: 0.1% ammonium bicarbonate in water; Mobile phase B: acetonitrile) to afford (*E*)-3-(7-(1-(2-(4-(2-(4-((4-([1,2,4]triazolo[1,5-a]pyridin-7-yloxy)-3-methylphenyl)amino)quinazolin-6-yl)vinyl)piperidin-1-yl)acetyl)piperidin-4-yl)-1-methyl-1*H*-indazol-3-yl)-1-methylpiperidine-2,6-dione (Tuc-CRBN, 622 mg, 696.5 µmol, 26.7% yield, 99.84% purity). LC-MS (ESI) *m*/*z* calculated for $C_{48}H_{49}N_{11}O_4$ $[M + H]^+$ 844.40, found: 844.10. $^1H$ NMR (400 MHz, DMSO-$d_6$): δ 10.89 (br s, 1H), 9.80 (br s, 1H), 8.95 (d, *J* = 7.60 Hz, 1H), 8.57 (s, 1H), 8.52 (s, 1H), 8.39 (s, 1H), 7.96 (d, *J* = 8.80 Hz, 1H), 7.91-7.84 (m, 2H), 7.72 (d, *J* = 8.40 Hz, 1H), 7.55 (d, *J* = 8.00 Hz, 1H), 7.25-7.18 (m, 2H), 7.12-7.02 (m, 2H), 6.81 (d, *J* = 2.40 Hz, 1H), 6.58-6.53 (m, 2H), 4.62-4.54 (m, 1H), 4.38-4.27 (m, 2H), 4.26 (s, 3H), 3.85-3.54 (m, 6H), 3.25-3.15 (m, 3H), 3.12-2.85 (m, 2H), 2.66-2.52 (m, 1H), 2.23 (s, 3H), 2.20-2.12 (m, 2H), 2.11-1.92 (m, 5H), 1.92-1.61 (m, 3H), 1.69.155 (m, 1H) ppm.

*Tuc-CRBN [1]H NMR (DMSO-d₆)*

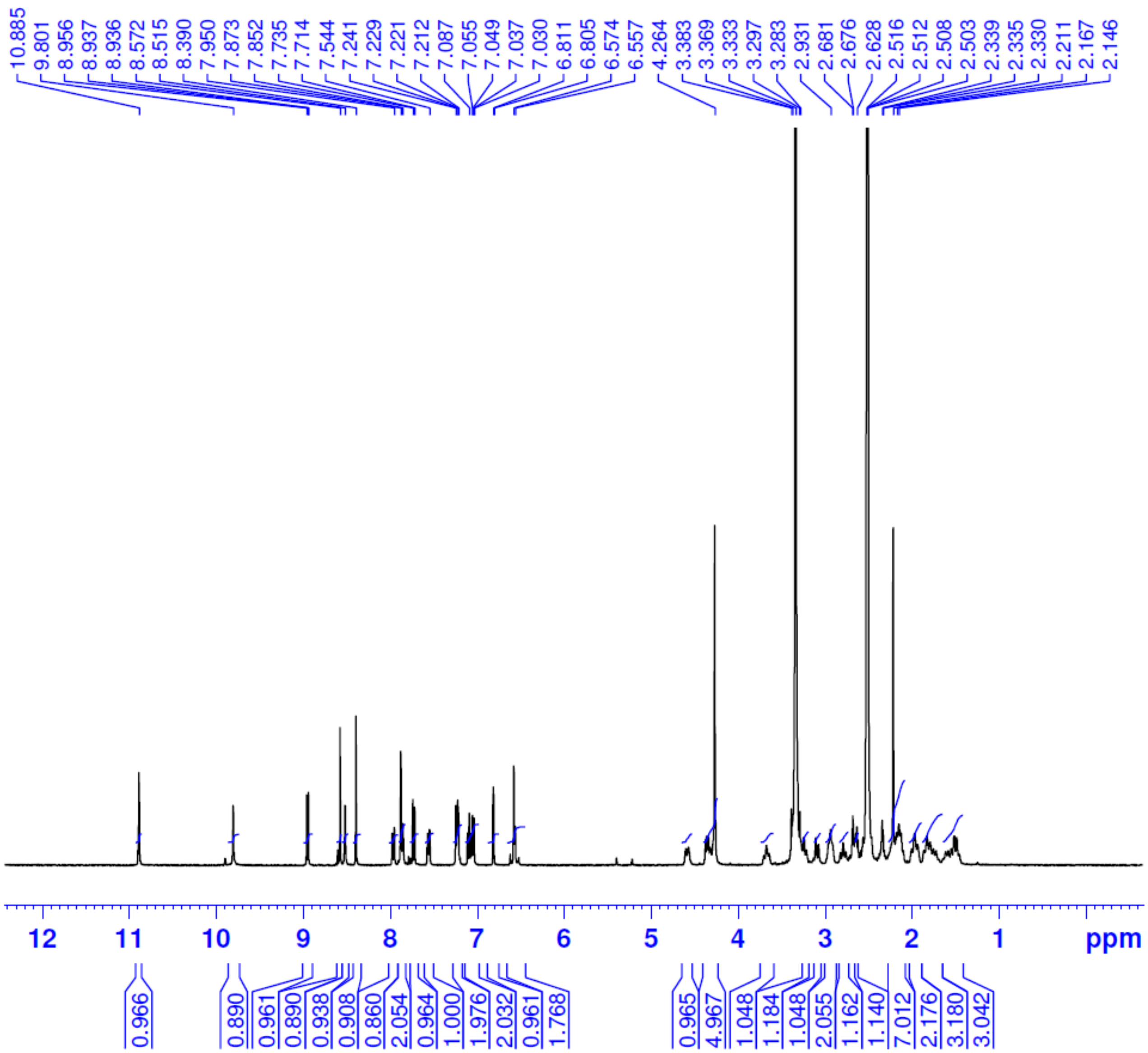

*Tuc-CRBN HPLC Purity Determination*

#### <Method Information>

Column:X-Select C18 (4.6X150mm,5μm)
MobilephaseA : 0.1%TFA in H20
MobilephaseB : ACN
Flowrate:2.0ml/min

| Time(min) | %B |
|---|---|
| 0.01 | 5 |
| 8.00 | 100 |
| 8.01 | 5 |
| 10.0 | Stoptime |

#### <Chromatogram>

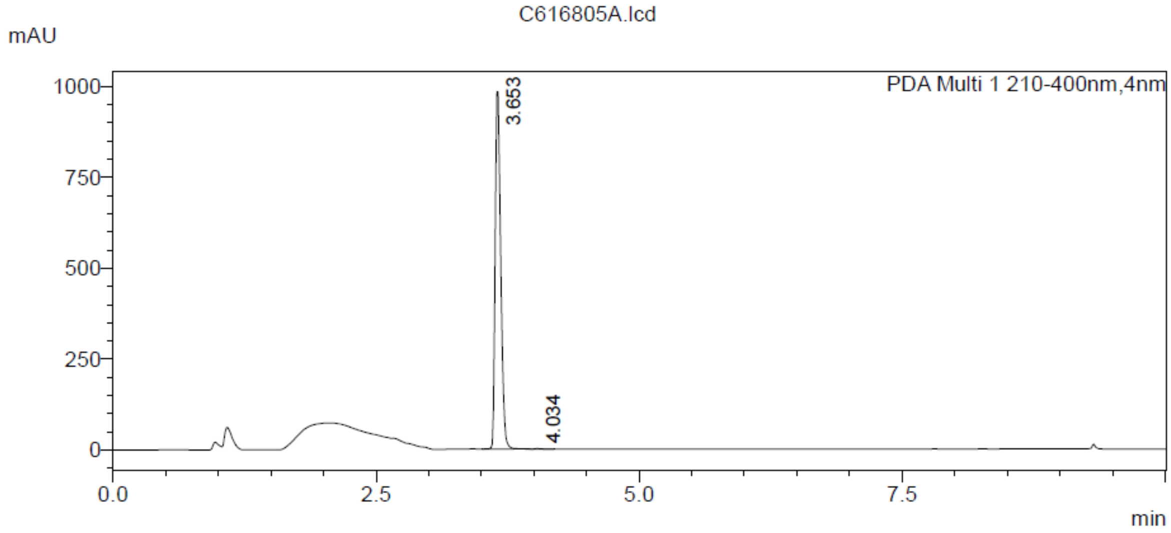


#### <Peak Table>

C616805A.lcd

PDA Ch1 210-400nm

| Peak# | Ret. Time | Height | Area | Area% |
|---|---|---|---|---|
| 1 | 3.653 | 983074 | 3640942 | 99.842 |
| 2 | 4.034 | 1403 | 5764 | 0.158 |
| Total | | 984478 | 3646707 | 100.000 |

*Synthesis of Intermediate 14*

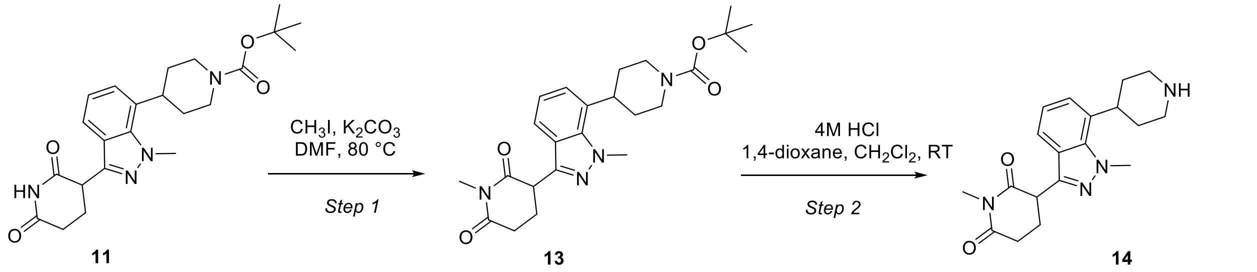


Step-1. To a solution of *tert*-butyl 4-[3-(2,6-dioxo-3-piperidyl)-1-methyl-indazol-7-yl]piperidine-1-carboxylate 11 (200 mg, 468.92 µmol, 1.0 eq) in DMF (3 mL) was added iodomethane (133.12 mg, 937.85 umol, 58.39 uL, 2.0 eq), and the resultant mixture was heated to 80° C for 3 h. The reaction mixture was diluted with ethyl acetate and water, extracted with EtOAc (2 x 15mL), and washed with brine (15 mL). The combined organics were dried over $Na_2SO_4$, filtered, and concentrated under reduced pressure to afford *tert*-butyl 4-[1-methyl-3-(1-methyl-2,6-dioxo-3-piperidyl)indazol-7-yl]piperidine-1-carboxylate 13 (200 mg, 408.60 umol, 87.13% yield, 90% purity) as a brown solid. LCMS (ES+) *m/z* calculated for $C_{24}H_{32}N_4O_4$ $[M+H]^+$ 441.25, found 441.30.

Step-2. To stirred solution of *tert*-butyl 4-[1-methyl-3-(1-methyl-2,6-dioxo-3-piperidyl)indazol-7-yl]piperidine-1-carboxylate 13 (200 mg, 453.99 umol, 1.0 eq) in dichloromethane (5 mL) was added trifluoroacetic acid (1.04 g, 9.08 mmol, 699.54 uL, 20 eq) at 0°C. The reaction mixture was stirred at ambient temperature for 3 hours, then concentrated under reduced pressure. The resultant residue was azeotroped with dichloromethane (2 x 10 mL) and triturated with methyl *tert*-butyl ether to afford 1-methyl-3-[1-methyl-7-(4-piperidyl)indazol-3-yl]piperidine-2,6-dione 14 (200 mg, 426.90 umol, 94.03% yield, 97% purity) as a viscous liquid/gum. LCMS (ES+) *m/z* calculated for $C_{19}H_{24}N_4O_2$ $[M+H]^+$ 341.20, found 341.20.

*Synthesis of Tuc-CRBN$_{Inactive}$*

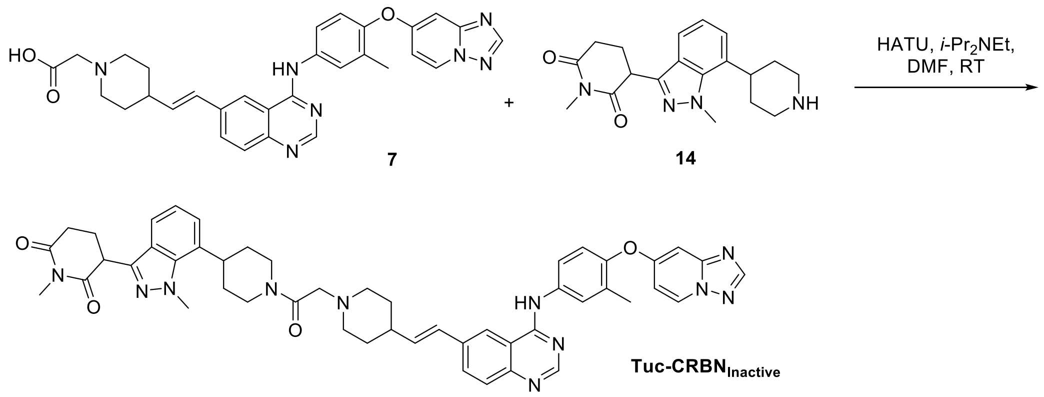

To a solution of 2-[4-[(*E*)-2-[4-[3-methyl-4-([1,2,4]triazolo[1,5-a]pyridin-7-yloxy)anilino]quin-azolin-6-yl]vinyl]-1-piperidyl]acetic acid 7 (80 mg, 139.9 µmol, 1.0 eq) and 1-methyl-3-[1-methyl-7-(4-piperidyl)indazol-3-yl]piperidine-2,6-dione 14 (63.6 mg, 139.9 µmol, 1.0 eq) in DMF (1.5 mL) was added *N*,*N*-diisopropylethylamine (122 µL 699.2 µmol, 5.0 eq) followed by HATU (79.8 mg, 209.8 µmol, 1.5 eq) under nitrogen atmosphere. The resultant reaction mixture was stirred at room temperature for 3 h. The reaction mixture was quenched with ice-cold water (5 mL) and stirred for 10 minutes. The precipitated solid was filtered and dried in vacuo to afford the crude compound, which was purified by reverse phase column chromatography (Column: Redisep $R_f$ Gold C18 100 g, Mobile phase A: 0.1% formic acid in water; Mobile phase B: acetonitrile) to afford (*E*)-3-(7-(1-(2-(4-(2-(4-((4-([1,2,4]triazolo[1,5-a]pyridin-7-yloxy)-3-methylphenyl)amino)quinazolin-6-yl)vinyl)piperidin-1-yl)acetyl)piperidin-4-yl)-1-methyl-1*H*-indazol-3-yl)-1-methylpiperidine-2,6-dione (Tuc-CRBN$_{Inactive}$, 29 mg, 31.88 µmol, 22.8 % yield, 99.37% purity. LC-MS (ESI) *m*/*z* calculated for $C_{49}H_{51}N_{11}O_4$ $[M + H]^+$ 858.40, found: 858.10. $^1H$ NMR (400 MHz, DMSO-$d_6$): δ 8.92 (d, *J* = 7.60 Hz, 1H), 8.58 (s, 1H), 8.55 (s, 1H), 8.38 (s, 1H), 8.22 (s, 1H), 7.95 (d, *J* = 8.80 Hz, 1H), 7.88-7.76 (m, 2H), 7.72 (d, *J* = 8.80 Hz, 1H), 7.56-7.53 (m, 1H), 7.23-7.18 (m, 2H), 7.10-7.03 (m, 2H), 6.78 (d, *J* = 2.40 Hz, 1H), 6.61-6.50 (m, 1H), 4.58-4.55 (m, 1H), 4.47-4.43 (m, 1H), 4.28-5.25 (m, 4H), 3.60-3.64 (m, 3H), 3.30-3.16 (m, 2H), 3.15-3.09 (m, 2H), 3.04-3.01 (m, 4H), 2.99-2.87 (m, 2H), 2.82-2.70 (m, 3H), 2.30-2.10 (m, 6H), 1.99-1.92 (m, 2H), 1.88-1.65 (m, 3H), 1.65-1.40 (m, 3H).

*Tuc- CRBN$_{Inactive}$ $^{1}$H NMR (DMSO-$d_6$)*

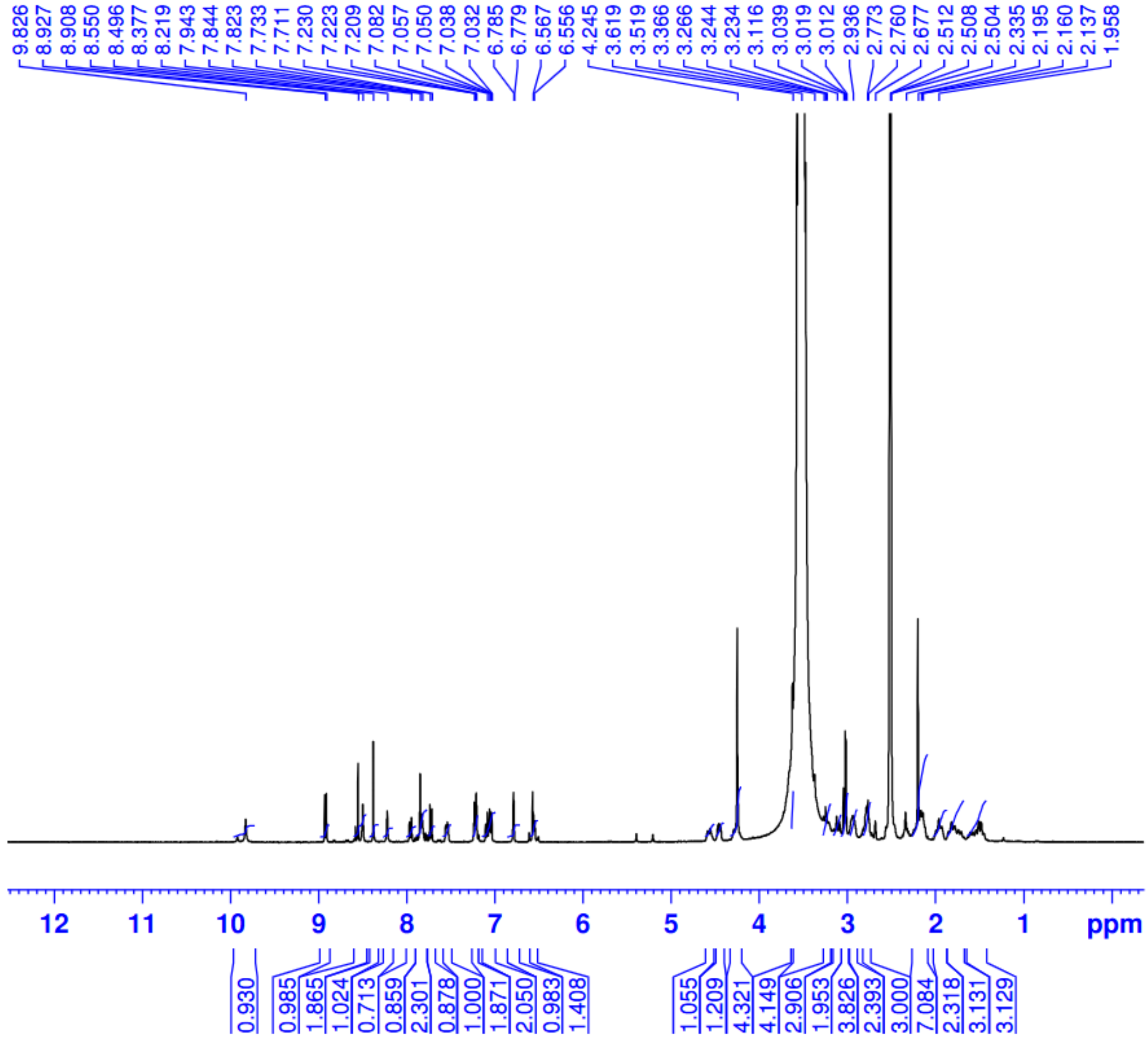

*Tuc-CRBN$_{Inactive}$ LC-MS Purity Determination*

### <Method Information>

Column :Acquity uplc BEH C18(2.1x50)mm,1.7µ
Mobile phase A:10mM Ammonium Bicarbonate in water
Mobile phase B:ACN
Flow rate :0.60 mL/min.

| Time(min) | %B |
|---|---|
| 0.01 | 5 |
| 0.30 | 5 |
| 2.50 | 100 |
| 3.70 | 100 |
| 3.80 | 5 |
| 4.00 | 5 |

#### <Chromatogram>

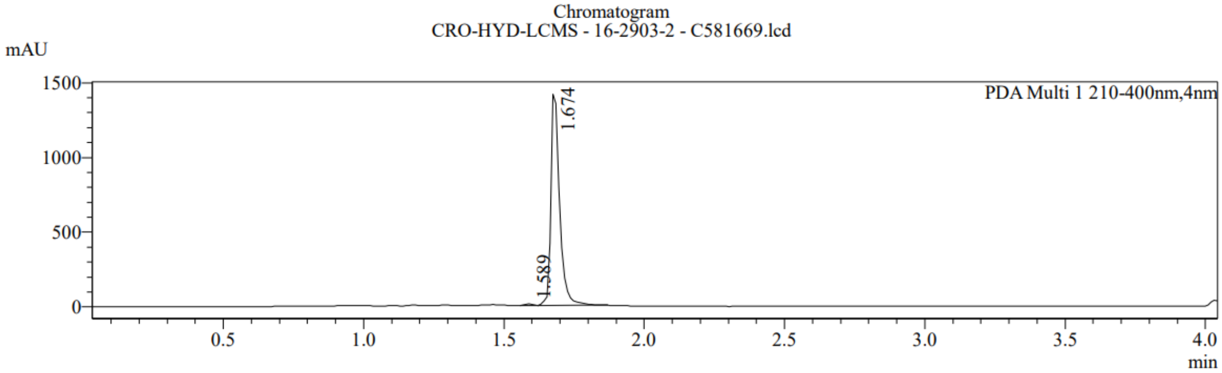


Peak Table C581669.lcd

PDA Ch1 210-400nm

| Peak# | Ret. Time | Area | Area% |
|---|---|---|---|
| 1 | 1.589 | 19768 | 0.627 |
| 2 | 1.674 | 3131214 | 99.373 |
| Total | | 3150981 | 100.000 |
